# Multilayer modelling and analysis of the human transcriptome

**DOI:** 10.1101/2020.05.21.109082

**Authors:** Tiago Azevedo, Giovanna Maria Dimitri, Pietro Lio, Eric R. Gamazon

## Abstract

Here, we performed a comprehensive intra-tissue and inter-tissue network analysis of the human transcriptome. We generated an atlas of communities in co-expression networks in 49 tissues (GTEx v8), evaluated their tissue specificity, and investigated their methodological implications. UMAP embeddings of gene expression from the communities (representing nearly 18% of all genes) robustly identified biologically-meaningful clusters. Methodologically, integration of the communities into a transcriptome-wide association study of C-reactive protein (CRP) in 361,194 individuals in the UK Biobank identified genetically-determined expression changes associated with CRP and led to considerably improved performance. Furthermore, a deep learning framework applied to the communities in nearly 11,000 tumours profiled by The Cancer Genome Atlas across 33 different cancer types learned biologically-meaningful latent spaces, representing metastasis (*p* < 2.2 × 10^−16^) and stemness (*p* < 2.2 × 10^−16^). Our study provides a rich genomic resource to catalyse research into inter-tissue regulatory mechanisms and their downstream phenotypic consequences.

## Introduction

The modern science of networks has contributed to notable advances in a range of disciplines, facilitating complex representations of biological, social, and technological systems (*1*). A key aspect of such systems is the existence of *community structures*, wherein groups of nodes are organised into dense internal connections with sparser connections between groups. Community structure detection in genome-wide gene expression data may enable detection of regulatory relationships between regulators (e.g., transcription factors or microRNAs) and their targets and capture novel tissue biology otherwise difficult to reach. Furthermore, it offers opportunities for data-driven discovery and functional annotation of biological pathways.

We hypothesise that community structure is an important organising principle of the human transcriptome, with critical implications for biological discovery and clinical application. Co-expression networks, in fact, encode functionally relevant relationships between genes, including gene interactions and coordinated transcriptional regulation, and provide an approach to elucidating the molecular basis of disease traits. Therefore, reconstructing communities of genes in the transcriptome may uncover novel relationships between genes, facilitate insights into regulatory processes, and improve the mapping of the human diseasome.

In this work, we develop a model of the human transcriptome as a multilayer network, and we perform a comprehensive analysis of the communities obtained with this modelling in order to further our understanding of its wiring diagram. We conduct a systematic analysis of the tissue or cell-type specificity of the communities in the transcriptome in order to gain insights into gene function in the genome and improve our ability to identify disease-associated genes. Our study represents an effort to fill an important gap in our understanding of the role of gene expression in complex traits, i.e., how a gene’s phenotypic consequence on disease or trait (*2*) is mediated by its membership in tissue-specific biological modules as molecular substrates. Methodologically, we demonstrate an approach to integrating the communities into transcriptome-wide association studies (TWAS) (*3, 4*) and a deep neural network methodology for generating biologically-meaningful latent representations of gene expression (*5, 6*). Finally, the inter-tissue analysis of the transcriptome that we present holds promise for identifying novel regulatory mechanisms and enhancing our understanding of trait variation and pleiotropy.

## Results

### Spurious Co-expression and Confounding due to Unmodelled Factors

Disambiguating true co-expression from artefacts is an important concern in the presence of hidden variables. We therefore applied *sva* analysis to investigate unmodelled and unmeasured sources of expression heterogeneity. The number of factors or components identified by this analysis was significantly correlated (*r* ≈ 0.95, *p* ≈ 5.4 × 10^−26^) with the number of samples across tissues (see Figure 1a). Notably, the greater number of such surrogate variables that we regressed out for tissues with larger sample sizes recapitulates the approach used by the GTEx Consortium (using a related adjustment method, i.e., PEER) of using more inferred factors for tissues with larger sample sizes (i.e., from 15 factors for tissue sample size *N* < 150 to 60 factors for *N* ≥ 350) in order to optimise the number of eGenes from the eQTL analysis (*7*).

**Figure 1:**
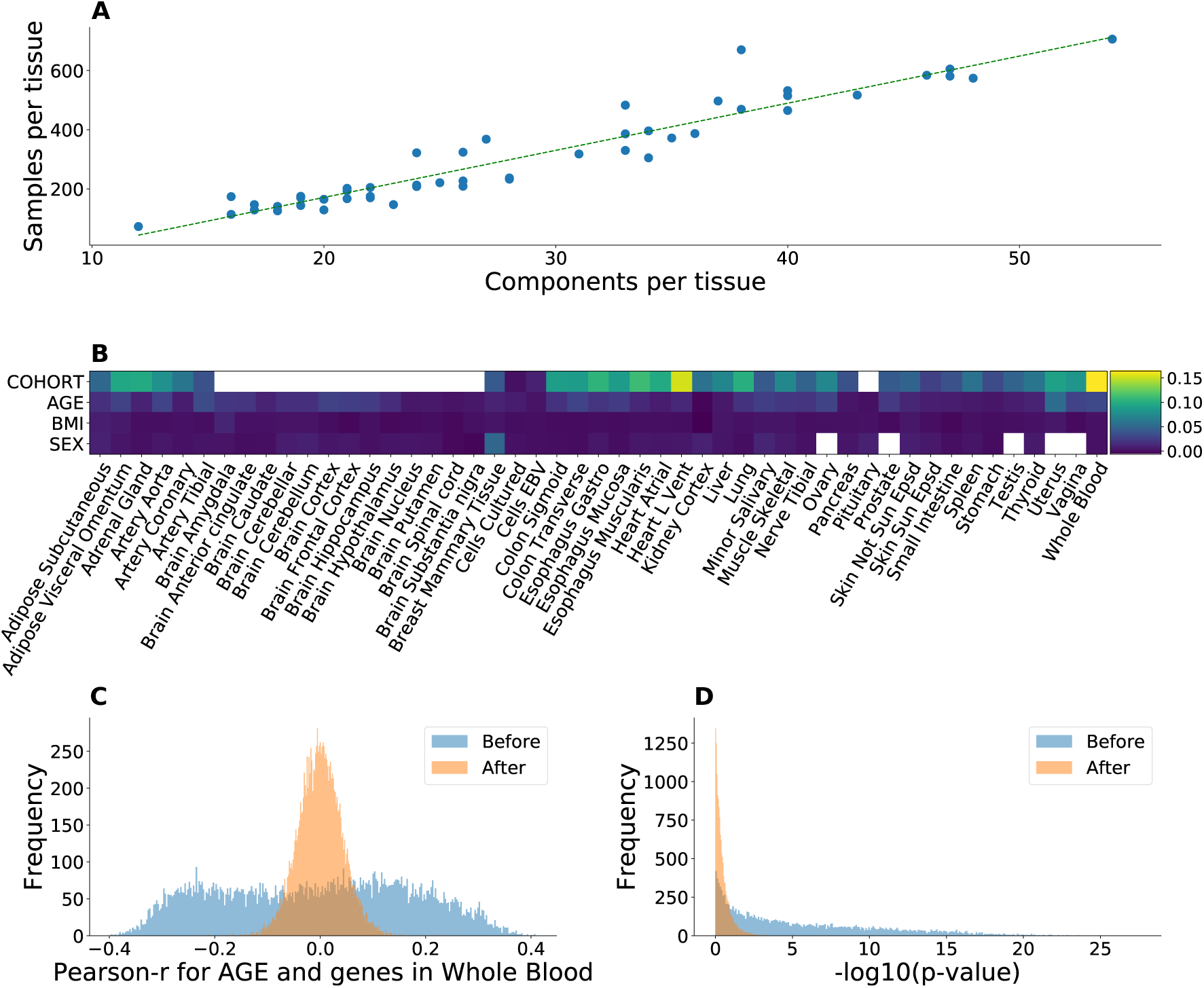
Confounding due to unmodelled factors. **(A)** Relationship between the number of inferred factors and tissue sample size. Fitted line (*r* ≈ 0.95, *p* ≈ 5.4 ×10^−26^) corresponds to a linear least-squares regression. The two-sided p-value is based on the null hypothesis that the slope is zero, using the Wald Test with t-distribution for the test statistic. **(B)** The difference in the variance of the distribution of Pearson correlation values for each tissue over all genes, before and after correction. Empty cells correspond to tissues in which only one value of the confound is available. The “Cohort” variable undergoes the most substantial change after the correction across all tissues. **(C)** Distribution of Pearson correlation between the expression of a gene in whole blood and age, before and after correction. After the correction, the correlation values move towards zero and show considerably less dispersion. **(D)** The p-value distribution from Panel (A)’s values, in logarithmic space. The enrichment for significant (low) p-values is greatly attenuated after the correction, suggesting that unmeasured variables can induce spuriously significant correlations.

We then quantified the impact of confound correction (see Figure 1b) in co-expression analysis. The distribution of correlation (Pearson) values is closer to zero with less variance after correction, suggesting that unmodelled factors may induce spurious (or artificially inflate) correlations in gene expression. The effect of unmodelled factors is further illustrated in Figure 1, Panels (B) and (C), where the distribution of correlation values for the covariate *Age* is shown for whole blood. Before correction, those values are spread between around −0.4 and 0.4, whereas after correction the corresponding values move towards the centre (zero) and become less dispersed. Notably, the variable *Cohort* (with possible values being *Postmortem* and *Organ Donor* in available tissues, except for some which also have *Surgical* values) seems to have undergone the largest change in the correction process. This suggests that estimation of cohort effect on gene expression can be substantially improved by accounting for unmodelled factors.

### Atlas of Communities across Human Tissues

For each tissue, we identified communities in the co-expression networks, using the Louvain algorithm (see Methods), to develop an atlas across human tissues. On average, a tissue was found to have 108 communities (standard deviation [SD] = 31) (see Figure 2). We observed the highest number of communities (*n* = 251) in “Kidney Cortex” and the lowest number (*n* = 73) in “Muscle Skeletal”. The nonsolid tissues, consisting of “Cells EBV” and “Whole Blood”, have the highest number of genes (i.e., at least 4,300 for each) that belong to a community. The size of a community varies considerably within each tissue and its distribution differs across tissues. (See Table S2 and Figure S2 for the distribution in all tissues.) The brain tissues (median SD = 9.9) show significantly higher variability (Mann-Whitney U test *p* = 1.55 × 10^−4^) than non-brain tissues (median SD = 5.18). Thus, tissues and tissue classes may differ in the overall topology of the communities in co-expression networks, which likely contains a lot of tissue information.

**Figure 2:**
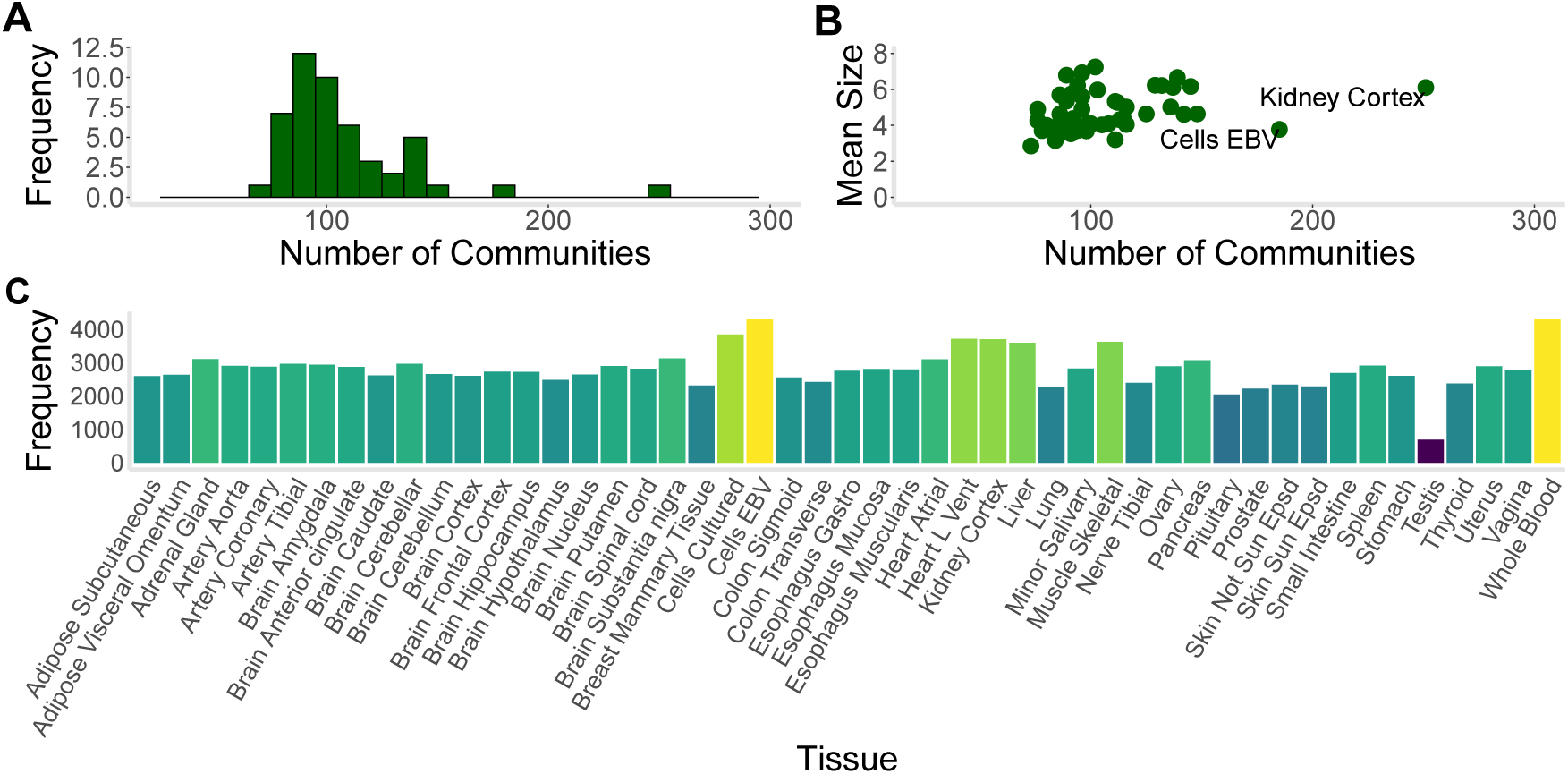
Summary statistics on identified communities. **(A)** Histogram shows the distribution of community count in the various tissues (mean = 108, SD = 31). **(B)** The scatter plot displays the community count and mean community size for each tissue, showing a significant correlation (Spearman *ρ* = 0.39, *p* = 0.006). The highest number of communities was observed in “Kidney Cortex” (*n* = 251). **(C)** Plot provides the number of genes that belong to a community in each tissue. The nonsolid tissues, “Cells EBV” and “Whole Blood”, show the highest number of genes with membership in a community.

We noticed that after removing the weaker correlations (−0.80 < *z*_*ij*_ < 0.80), most of the subnetworks were already highly segregated from the rest of the entire network, indicating that the Louvain communities could almost be completely formed by just this removal process. In order to evaluate the segregation of such communities, we calculated the number of connections coming out of communities of each size. We found that for every tissue the mode was zero, and the maximum number was never over 17. Given the thousands of genes in each tissue’s co-expression network, the observed maximum number of connections between different communities (i.e., at most 17) illustrates how strong the segregation is prior to the application of the Louvain community analysis.

More information on these communities is available on github: notebook *09 community info*.

### UMAP of Community-defined Gene Expression Manifold Reveals Tissue Clusters

To generate a lower dimensional representation of the original transcriptome dataset, we performed Uniform Manifold Approximation and Projection (UMAP) (*8*) (see Methods). Nearly 18% of the genes belong to a community in at least one tissue. Notably, gene expression from this subset was able to recover the tissue clusters (see Figure 3a) as fully as the complete set of genes analysed here (see Figure S1).

**Figure 3:**
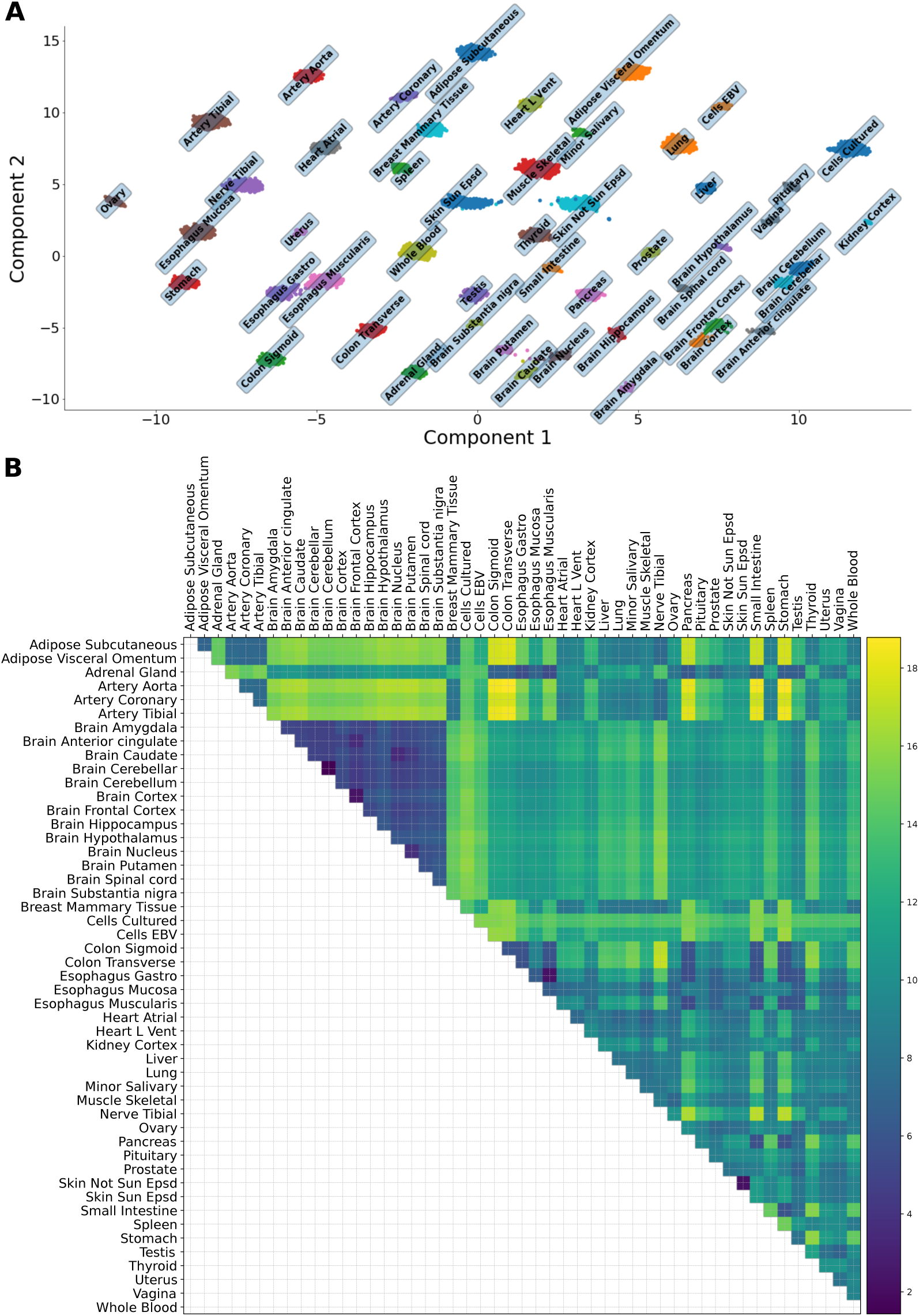
Lower-dimensional UMAP representation of the transcriptome data restricted to the communities and conservation of global structure. **(A)** UMAP generates embedded structures through a low-dimensional projection of the submatrix consisting of only the genes (*n* = 3, 259) that belong to a community in at least one tissue. This subset of genes (17.7% of total) contains sufficient information to recover the tissue clusters. In addition, known relationships between tissues, based on organ membership and, separately, on shared function, are reflected in the UMAP projection. **(B)** Using bootstrapped manifolds (see Methods), we estimated the persistence of the global structure and pairwise relationships across tissue clusters. Here we show the upper-triangular matrix of the average pairwise distances across the bootstrapped manifolds. We found consistent clustering of known related tissues, including the 13 brain regions, the colonic and esophageal tissues, and various artery tissues. Additional patterns were observed. For example, as reflected in the heatmap, we found a highly correlated relationship, i.e., high “clustering conservation coefficient” 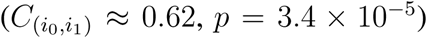 (see Methods), of the two skin tissues to all the other tissues

Drawing conclusions about relationships between clusters (tissues) from UMAP (and related approaches) must be done with caution due to some known caveats (*9*) (see next section for more details). However, starting from known relationships between tissues, we found that the subset of community-based genes yielded biologically consistent embeddings from UMAP. Indeed, the clustering of related tissues (based on organ membership), such as the 13 brains regions, or the clustering of other related tissues (based on shared function), such as the hypothalamus-pituitary complex (which controls the endocrine system (*10*)), could be observed for the genes that belong to communities. Taken together, these results show that gene expression from the identified communities encodes sufficient information to distinguish the various tissues in a biologically-meaningful low-dimensional representation. We note, however, that not all sets of genes with correlated expression produce the distinct separation of tissues observed for the set of genes that belong to the communities (see below).

In theory, additional clusters may be present at different scales, e.g., within a tissue. Therefore, we performed UMAP analysis on the single-tissue “Whole Blood” to test for the presence of additional clusters. Notably, no well-defined clustering was observed with respect to cohort (Figure S3), BMI (Figure S4), and the other covariates, indicating that the *sva* analysis was successful in removing potential confounders (see Figure 1b).

External transcriptome data can be embedded into the trained model generated from the GTEx communities. Indeed, using TCGA data for 3 cancer types, we found that the embeddings into the learned space recapitulate recent findings on cancer-testis (CT) genes (see Supplementary Material).

### Persistence of the UMAP Embeddings

We developed an approach to quantify the conservation and variability of the UMAP global structure and estimate the sampling distribution of the local structure, e.g., the distance d(*i, j*) for a given pair of tissues *i* and *j* (see Methods). Using 500 bootstrapped manifolds, we found, for example, that, on average, (a) the 13 brain regions, (b) the colonic and esophageal tissues, and (c) various artery tissues tended to cluster closely together (see Figure 3b). Figure S5 shows the relationship between the average distance between tissue clusters and the variance in the distance, showing a significant positive correlation (Spearman *ρ* ≈ 0.38, *p* < 2.2 × 10^−16^). Reassuringly, the tissue pairs (“Brain Cerebellum”, “Brain Cerebellar”) and (“Skin Not Sun Epsd”, “Skin Sun Epsd”) had the lowest average distance between clusters among all tissue pairs; the first pair consists of known duplicates of a brain region in the GTEx data (*11*) and is thus expected to cluster together. Among the tissue pairs with the highest average distance, “Adipose Subcutaneous” with each of the colonic tissues (“Colon Sigmoid” and “Colon Transverse”, each with average distance greater than 17) had low variance comparable to tissue pairs with some of the smallest average distance. Additional global patterns can be easily observed.

For example, the relationships of related tissues (e.g., “Skin Sun Epsd” and “Skin Not Sun Epsd” with 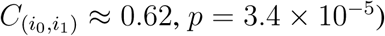 to the remaining tissues were found to be strongly preserved, using our clustering conservation coefficient (see Methods).

### Prediction of Tissues by Communities

We then tested individual communities for their ability to predict a tissue. By definition, we consider that a set of genes can predict a tissue when the average *F*_1_ score is above 0.80 (see Methods). Some broad patterns are noteworthy. Most of the communities from “Whole Blood” do not have prediction power for the other tissues (Figure 4a) partly due to the stringency of our *F*_1_ threshold, which is likely to produce false negatives. This observation indicates that the member genes in each such community from the source tissue (“Whole Blood”) cannot “separate” the test tissue (say, “Lung”) from the remaining tissues possibly due to lack of tissue specificity of the gene expression profile of the community. (Here we tested a linear classifier, and the so-called “kernel trick” may work in the non-linearly separable gene expression profiles, though perhaps at the expense of biological interpretability.) However, a community of only five genes can predict the brain region nucleus accumbens (basal ganglia). For these communities, the member genes, collectively, are “differentially expressed” between the test brain region and the remaining tissues. Thus, although the genes are present in “Whole Blood” (as a community), the expression profile in the test brain region is substantially different or tissue-specific. “Cells cultured fibroblasts” is the tissue which can be predicted by the largest number of “Whole Blood” communities (three) and, consistently, the largest number from the other tissues.

**Figure 4:**
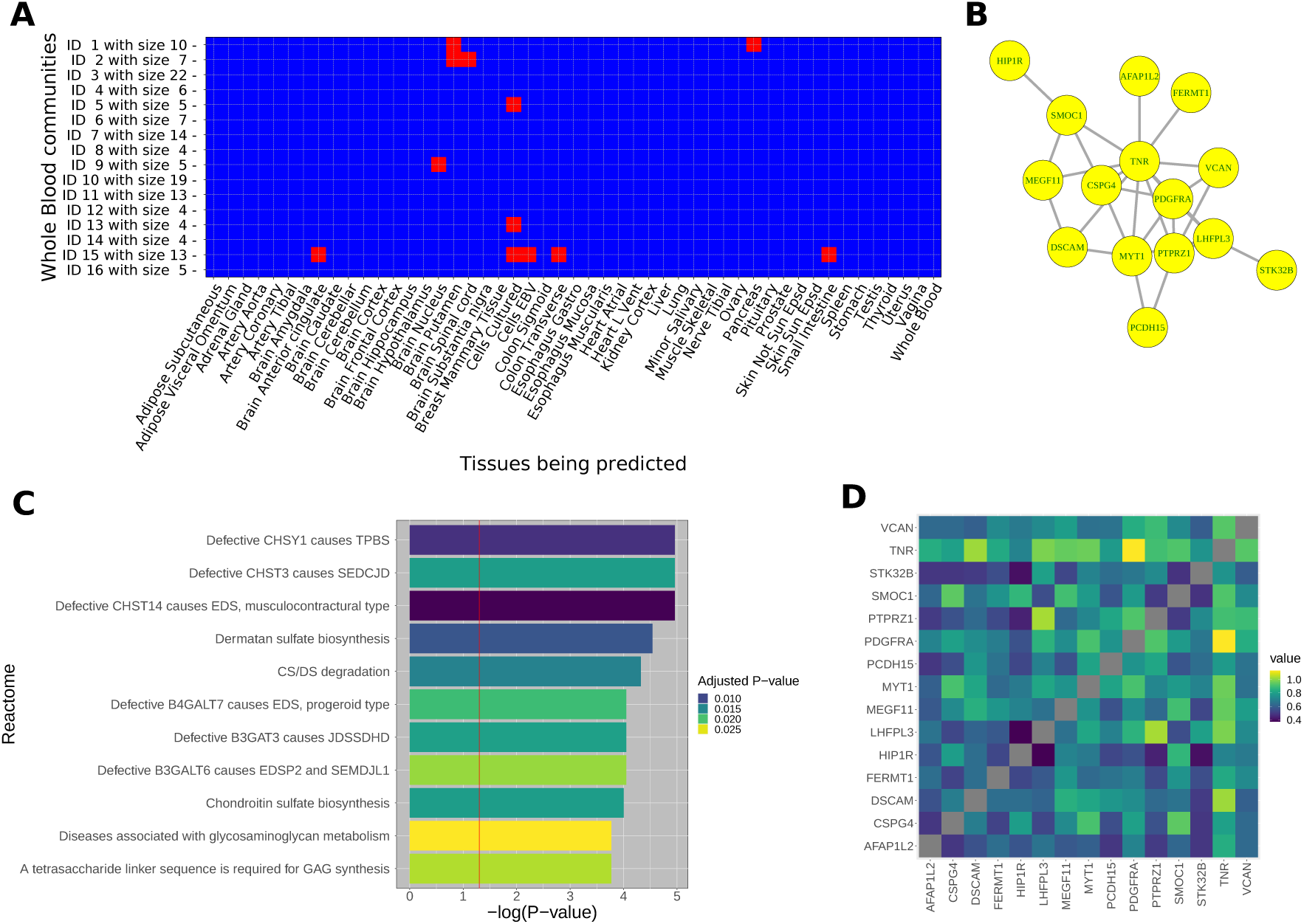
Communities and their properties. We developed tools (available on github) to query a community for its characterisation. **(A)** Prediction power of “Whole Blood” communities, in F1 scores thresholded over 0.8. Most communities in “Whole Blood” do not have prediction power for the remaining tissues. Notable exceptions include a 13-member community, which can predict multiple tissues, including “Brain Anterior cingulate”, “Small Intestine”, and “Colon Transverse”. **(B)** A 15-member community in the hippocampus is shown here as an example. An edge indicates *A*_*ij*_ > 0.80 for genes *i* and *j*. The gene *TNR*, which is expressed primarily in the central nervous system and involved in its development, is connected via an edge to twelve member genes while *HIP1R* is connected to only one. **(C)** Enrichment analysis was performed on all communities to identify known biological processes. For example, the hippocampal community in panel (B) was found to be significantly enriched for Reactome pathways. P-value refers to raw p-value. Red line corresponds to the raw *p* < 0.05 threshold. Colour gradient reflects the adjusted p-value. All Reactome pathways shown meet adjusted *p* < 0.05. **(D)** Heatmap displays the correlation values for the member genes of the community in panel (B).

We note that, consistent with our observations for the communities, most Reactome pathways are not sufficient to predict any tissue (available on github: output *output 06 02*), while many are tissue-specific (i.e., can predict only one tissue) (see Supplementary Material).

### Enrichment of Communities for Known Biological Processes

We quantified the extent to which the communities in the various tissues reflect current biological knowledge (as encoded in the Reactome pathways). We identified 114 communities (8.28% of all the communities with more than 3 member genes) enriched for some Reactome pathway (i.e., at an adjusted *p* < 0.05 for level of enrichment), thus contributing in complex ways to multiple biomolecular processes. “Whole Blood” was the only tissue without any community enriched for known pathways, and the “Esophagus Mucosa” was the tissue with the most communities enriched for known pathways, with a total of 5 communities. Since the entire set of communities could fully recover all tissues as clusters in the UMAP embeddings, these results suggest that the remainder of the communities are likely to capture previously inaccessible and novel tissue biology.

Notably, our analysis may uncover the role of these communities in human diseases. For example, a community of 15 genes in the “Brain Hippocampus” showed a significant enrichment for diseases associated with glycosaminoglycan metabolism (adjusted p-value = 0.026; see Figure 4). Glycosaminoglycans, which are major extracellular matrix components whose interactions with tissue effectors can alter tissue integrity, have been shown to play a role in brain development (*12*), modulating neurite outgrowth and participating in synaptogenesis. Alterations of glycosaminoglycan structures from Alzheimer’s disease hippocampus have been implicated in impaired tissue homeostasis in the Alzheimers disease brain (*13*).

### Multiplex Analysis of the Transcriptome

We analysed five multiplex networks to model the various tissue interactions, of clear biological interest, in the GTEx dataset (see Methods). For each multiplex architecture, only the specific component tissues were used to construct the multiplex network, and consequently we calculated the global community index for each multiplex architecture, using only the component tissues of the multiplex network.

- **All Tissues**: Each layer represents one of the 49 tissues analysed. This architecture allows us to investigate gene communities that are shared across all tissues, with potentially universal function.
- **Brain Tissues**: The 13 layers correspond to the various brain regions. This architecture facilitates identification of communities that may play a functional role throughout the central nervous system (CNS).
- **Brain Tissues and Whole Blood**: This multiplex model consists of the 14 layers corresponding to these tissues. This architecture allows us to study brain-derived communities for which the easily accessible whole blood can serve as a proxy tissue.
- **Brain and Gastrointestinal Tissues**: The 16 layers correspond to the brain tissues and 3 gastrointestinal tissues. This architecture may provide insights into the gut-brain axis, which has attracted recent attention in the literature, e.g., in the study of neuropsychiatric processes and the interaction between the CNS and the enteric nervous system in neurological disorders (*14, 15*).
- **Non-Brain Tissues**: The 36 layers consist of all tissues outside the brain. This architecture may stimulate investigations into developmental and pathophysiological processes outside the CNS.

Multiplex analysis provides an inter-tissue framework for the analysis of high-dimensional molecular traits such as gene expression. The global multiplexity matrix was obtained for each of the five proposed architectures. We extracted from the global multiplexity matrices the groups of genes with the maximal global multiplexity index in the five architectures, i.e., the groups of genes that share a value of 49, 13, 14, 16, and 36 respectively, equal to the number of layers (tissues) in the respective architectures. Among these groups of genes with the highest global multiplexity index, we obtained the sub-clusters for each architecture, identifying the groups of genes that always appear in the same community across the various layers. Revealing the shared community structure across the layers improves our understanding of the functional and disease consequences of the clusters of genes. We investigated the biological pathways (Reactome) in which such subgroups were involved for each architecture. Our goal was to test the communities for enrichment for known biological pathways and therefore quantify the degree to which the communities capture current understanding of biological processes as encoded in the knowledge base.

We illustrate this approach here. (The complete results for all 5 architectures can be found on our github page in the jupyter notebook *11 multiplex enrichment*.*ipynb*.) In Figure 5a, we show an example of a multiplex network. In this case, the multiplex network is constructed from data in two brain tissues: “Brain Hippocampus” and “Brain Putamen” (basal ganglia). Each layer represents a tissue, and nodes are labelled as the genes to which they correspond. In the structure, we can see the presence of two communities (in this case, randomly drawn from the full set) formed by the groups of genes: {*FGA, ORM1, FGB*} and {*MYH11, ACAT2, MYL9*}. Corresponding genes are connected through the layers as shown, through interlayer connections. We note that, based on our analysis, these two communities are indeed part of the “Brain Tissues” multiplex architecture, present in all 13 component layers (brain regions). Notably, all three genes that belong to the first community have been previously implicated as biomarker and therapeutic target candidates for intracerebral hemorrhage (*16*). This observation is interesting given our Reactome pathway analysis results; the community is enriched for “Common Pathway of Fibrin Clot Formation” (adjusted *p* = 8.8 × 10^−4^) and “Formation of Fibrin Clot (Clotting Cascade)” (adjusted *p* = 1.7 × 10^−3^), indicating the genes’ involvement in coagulation. All three members of the second community are myelin-associated genes (*17*). We found a 17-member community in the “Brain Tissues” multiplex that is significantly enriched for the nonsense-mediated decay (NMD) pathway (adjusted *p* = 1.01 × 10^−37^), which is known to be a critical modulator of neural development and function (*18*). The pathway accelerates the degradation of mRNAs with premature termination codons, limiting the expression of the truncated proteins with potentially deleterious effects. The community’s presence in all brain regions underscores its crucial protective function throughout the central nervous system.

**Figure 5:**
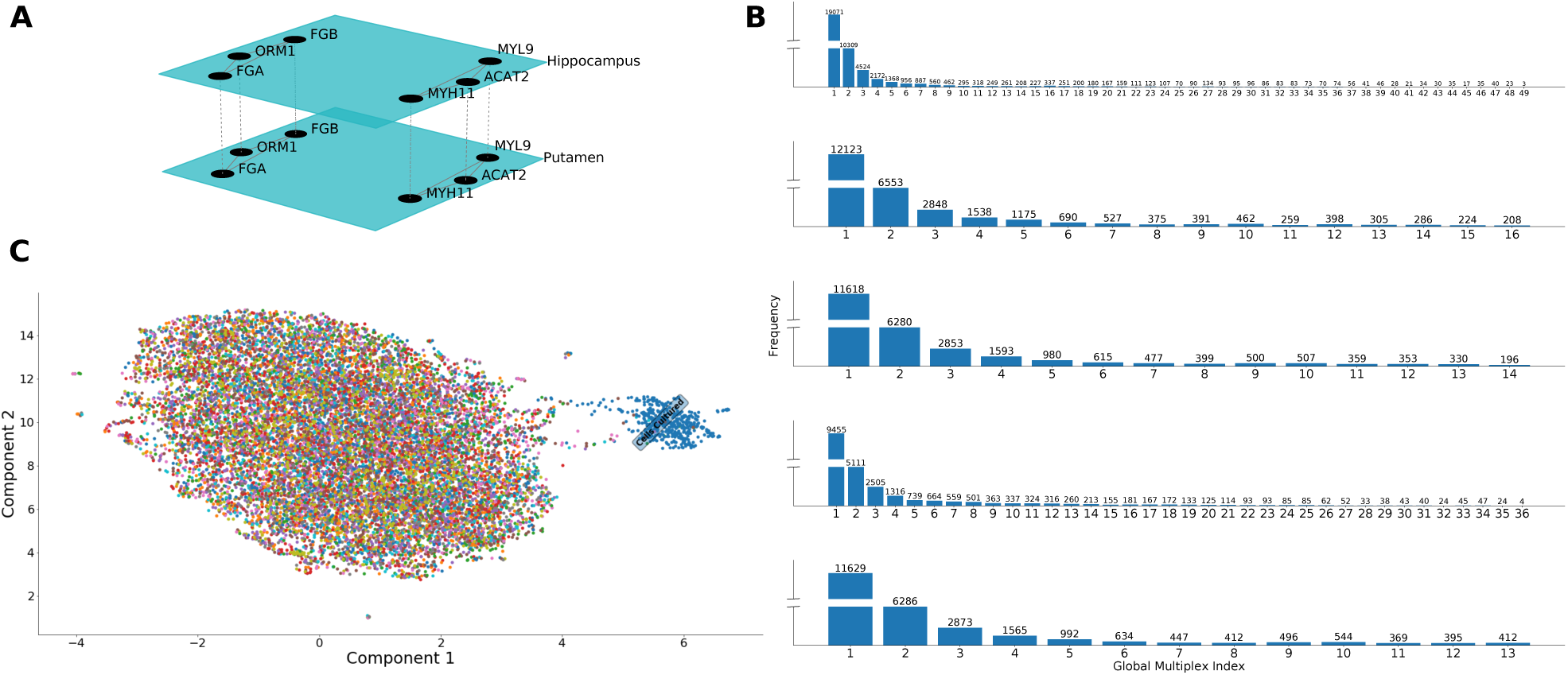
Multiplex analysis. **(A)** An example of a multiplex network in “Brain Hippocampus” and “Brain Putamen” (basal ganglia). Each layer denotes a tissue, and nodes are the genes, which are connected via the interlayer connections. In the multiplex example, we see the presence of two communities. These two communities are indeed part of the “Brain Tissues” multiplex architecture, present in all 13 layers (brain regions). All genes in the community {*FGA, ORM1, FGB*} have been implicated as biomarker and therapeutic targets for intracerebral hemorrhage. **(B)** Histograms show the empirical distribution of the global multiplexity index for each multiplex architecture (with positive index). The index quantifies how many times two genes belong to the same communities across layers. The proportion at each value *k* of the index is an estimate of the *π*_*k*_ (see Methods). The maximum value corresponds to the number of layers or tissues of the multiplex network. Histograms, from the top, correspond to: All tissues; Brain tissues; Brain tissues and whole blood; Brain and gastrointestinal tissues; Non-brain tissues. **(C)** We performed UMAP on the subset of communities that exist across all layers of the central nervous system (“Brain Tissues”) multiplex. This set does not yield complete clustering of tissues.

The multiplex analysis we performed can also be used to investigate the relationship between two distinct systems. Here we illustrate this using the CNS and the gastrointestinal system, possibly reflecting a coordinated transcriptional regulatory mechanism between the CNS and the enteric nervous system (ENS). The ENS is a large part of the autonomic nervous system that can control gastrointestinal behaviour (*19*). We found a 14-member community in the “Brain and Gastrointestinal Tissues” multiplex, whose presence in all 16 layers suggests a strong interaction between the CNS and ENS. Consistent with this hypothesis, the community was found to be significantly enriched for the “metabolism of vitamins and cofactors” (adjusted *p* = 6.5 × 10^−7^), which has been shown to be responsible for altered functioning of the CNS and ENS (*20*). Although the involvement of the individual member genes in this pathway is known, the finding that the genes are organised as a community structure, within co-expression networks, which persists across the entire 16 layers of the various brain regions and the gastrointestinal tissues examined here is a novel one.

The empirical distribution of the global multiplexity index is presented in Figure 5b for each of the five architectures. The maximal global multiplexity index in the five architectures represents the groups of genes that share a value of 49, 13, 14, 16, and 36 respectively, equal to the number of layers (i.e., tissues) in the respective architectures. These genes appear in the same community across *all* layers of the respective architectures.

For comparison with the UMAP embeddings of the set of all communities, we performed similar analyses in the various multiplex networks. For example, we tested whether the complete tissue clustering could be observed using just the subset of communities that exist across all layers of the central nervous system multiplex. We discovered a different clustering pattern, with cultured fibroblasts clustering separately from the rest of the tissues, which no longer show well-defined clustering (Figure 5c). This finding suggests the presence of a hierarchy of clusters in the transcriptome at increasingly finer scales.

### Communities and Transcriptome-Wide Association Studies of Disease

Leveraging the communities may have important methodological implications on the search for disease-associated genes. We asked whether incorporation of the communities would improve our ability to detect significant gene-level (TWAS / PrediXcan) associations (see Methods). We performed a TWAS of CRP in 361,194 UK Biobank subjects, a biomarker of chronic low-grade inflammation, with elevated CRP levels associated with a broad array of complex diseases, including cardiovascular disease, Alzheimer’s disease, and schizophrenia (*21*). Notably, we found a significantly greater enrichment for associations with CRP (defined as adjusted *p* < 0.05) among the set of genes that belong to a community than among the complement set of genes. In particular, the genes in communities showed a greater departure from the null (expected) distribution than the complement set of genes (Figure 6a). This observation suggests that the use of the communities can substantially improve the signal-to-noise ratio in TWAS even in the case where the dataset is already highly-powered to detect causal associations. The estimated true positive rate 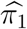 (see Methods) for association with the trait for the set of genes in communities was 0.45 while the estimate for the complement set was 0.37. Our top association with CRP among the community-located genes was *OASL* (*p* = 1.56 × 10^−55^; see Figure 6d for the chromosomal positions of the top associations), which has been previously implicated as a CRP and cardiovascular disease associated gene (*22*).

**Figure 6:**
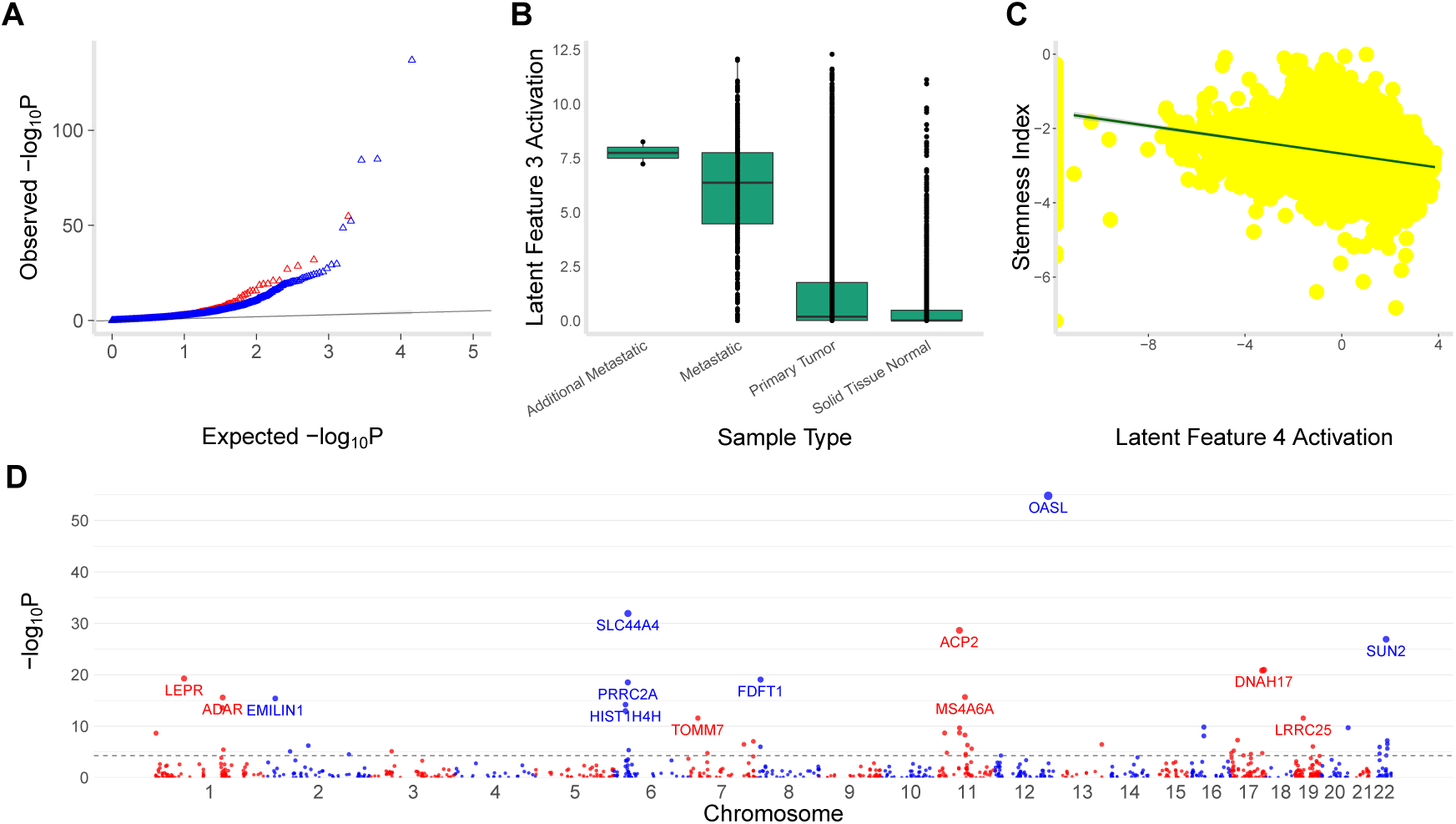
Integrating communities into TWAS and Variational Autoencoder. Methodological integration of the communities into genomic studies of disease may improve identification of disease-associated genes and enable unique biological insights. **(A)** We performed a transcriptome-wide association study (TWAS) of C-reactive Protein (CRP) in 361,194 UK Biobank individuals using PrediXcan. CRP is a biomarker for a wide range of complex diseases. The set of genes in the communities (red) displayed a greater departure from the null expectation (i.e., greater enrichment for significant associations) than the complement set of genes (blue), as shown by the leftward shift in the Q-Q plot. **(B)** We implemented a variational autoencoder (VAE) to leverage the power of neural networks in high-dimensional molecular data. Using 11,060 samples across 33 cancer types from TCGA, the VAE learned biologically-meaningful latent spaces from the communities. For example, latent feature 3 separates metastatic tumours from primary tumours and normal solid tissues (Mann-Whitney U test *p* < 2.2 ×10^−16^ for each comparison), representing a potential mechanism for metastasis. **(C)** The VAE model learned a stemness index significantly associated (*p* < 2.2 × 10^−16^) with a DNA-methylation-based index. **(D)** Manhattan plot shows TWAS associations with CRP for the genes in the communities (dashed line is Bonferroni-adjusted *p* < 0.05).

### Variational Autoencoder Model of Communities and Phenotypic Consequences

Methodologically, the communities may also enable discovery of biologically-meaningful features in high-dimensional molecular data. We implemented a variational autoencoder (VAE) model (see Methods) of the communities in 11,060 samples across 33 different cancer types in the TCGA data, customising the *Tybalt* approach (*5*). One benefit of a VAE is that it offers a probabilistic model, allowing us to do inference on the latent variable *z*, i.e., *P* (*z*|*X*). The marginal log-likelihood, log *P* (*X*), is generally intractable, which poses a challenge to this inference; however, equation (13) provides a variational lower bound on this marginal. In the VAE model, we assumed that the approximating posterior distribution *Q*(*z*|*X*) is multivariate Gaussian, whose mean *µ*(*X*) and diagonal covariance matrix σ^2^(*X*)**I** are learned by a neural network, so as to leverage the “reparametrisation trick” (*23*). The second stage, which is also implemented as a neural network, generates a reconstructed representation of *X* from the stochastic *z*. In our implementation, we randomly split the input into a training set (80%) and a test set (20%). The encoded layer is compressed into a vector of size 100 consisting of a mean and variance. We assumed a learning rate of 0.0005, 50 epochs, and batch size of 50. We used Rectified Linear Unit (ReLU) activation for the encoder and sigmoid activation for the decoder. The optimisation algorithm *Adam* was applied in the training to minimise the VAE loss, i.e., the negative of the Evidence Lower Bound Objective (ELBO) given by equation (13). The VAE loss as a function of epoch number (showing the training performance) and the reconstruction accuracy for the communities are largely equivalent to the corresponding results for the full transcriptome (Figure S6) in the TCGA data.

Notably, the latent representations learned by the VAE model from the communities encode biologically-meaningful features. The model learned to stratify metastatic tumours from primary tumours and normal solid tissues (Figure 6b) (Mann-Whitney U test *p* < 2.2×10^−16^). This held robustly after adjusting for ethnicity, sex, age, or stage (logistic regression *p* < 2.2 × 10^−16^ for each) or disease (*p* = 8.9 × 10^−5^). Cancer progression may be characterised by oncogenic dedifferentiation (i.e., steady loss of differentiated phenotype) and acquisition of stemness (i.e., self-renewal and generation of differentiated progeny). The VAE model learned a stemness index that was significantly associated (Spearman *ρ* ≈ −0.27 in log_2_ transformed space, *p* < 2.2 × 10^−16^), across the spectrum of tumour types, with a recently developed DNA-methylation based index (Figure 6c) (*24*) (see Methods). Again, adjustment for ethnicity, sex, age, stage, or disease did not affect the result (linear regression *p* < 2.2 × 10^−16^).

Finally, a freely available genomics resource of communities, their predictive power for tissues, and multiplex networks, and the software to construct a similar resource on a transcriptome dataset and to facilitate integration into genomic studies of disease can be accessed at https://github.com/tjiagoM/gtex-transcriptome-modelling.

## Discussion

We develop an inter-tissue multiplex framework for the analysis of the human transcriptome. Given the complexity of pathophysiological processes underlying complex diseases, intra-tissue and inter-tissue transcriptome analysis should enable a more complete mechanistic understanding. For these phenotypes, studying the interaction among tissues may provide greater insights into disease biology than an intra-tissue approach. Communities in co-expression networks are here shown to be enriched for some known pathways, encoding current understandings of biological processes; however, we identify other communities that are likely to contain novel or previously inaccessible functional information. Methodologically, integration of the communities into TWAS and a neural network demonstrate substantial gain in the identification of disease-associated genes and discovery of biologically-meaningful information.

UMAP embeddings of the transcriptome of the entire set of communities (representing only 18% of all genes) fully reveal the tissue clusters. Low-dimensional representation of the subset of communities that are in the multiplex networks does not recover the tissue clusters, but uncovers other clustering patterns, suggesting a hierarchy of clusters at increasingly finer scales. We develop an approach to quantify the conservation of, and uncertainty in, the UMAP global structure and estimate the sampling distribution of the local structure (e.g., distances among tissue clusters), with broad relevance to other applications (such as cell population identification in single-cell transcriptome studies). New gene expression data can be embedded into our models, facilitating integrative analyses of the large volume of transcriptome data that are increasingly available. We provide a publicly available resource of co-expression networks, communities, multiplex architectures, and enriched pathways, and code to stimulate research into network-based studies of the transcriptome.

Using the global multiplexity index, we investigate the tissue-sharedness of identified communities. In fact, communities that are shared across multiple tissues may suggest the presence of a tissue-to-tissue mechanism that controls the activity of member genes across the layers in the network. Such regulatory mechanisms have been relatively understudied in comparison with intra-tissue controls.

We identified tissue-dependent communities that are enriched for human diseases. For example, we found a 15-member community in the “Brain Hippocampus” that is enriched for diseases associated with glycosaminoglycan metabolism. These genetic disorders are due to mutations in the biosynt”hetic enzymes, e.g., glycosyltransferases and sulfotransferases, for glycosaminoglycans. Sulfated glycosaminoglycans include the chondroitin sulfate and dermatan sulfate chains that are covalently bound to the core proteins of proteoglycans, which are present in the extracellular matrices and at cell surfaces. Mutations affecting the biosynthesis of these chains may lead to genetic diseases that are characterized by craniofacial dysmorphism and developmental delay (*25*). The community structure we identified proposes a cooperative role for these genes, and the fact that they span multiple chromosomes suggests the presence of coordinated transcriptional regulation.

Some of the communities are shared across multiple tissues; their dysregulation may thus lead to pleiotropic effects and contribute to known and novel comorbidities. Here we modeled these communities as belonging to layers of multiplex networks. For example, we identified a 17-member community in the “Brain Tissues” multiplex network (i.e., spanning across all brain regions sampled here), consisting primarily of ribosomal proteins (therefore, enriched for proteins involved in translation [adjusted *p* = 2.53 × 10^−35^]), with significant overrepresentation for the viral mRNA translation pathway (adjusted *p* = 1.06 × 10^−38^), NMD (adjusted *p* = 1.01 × 10^−37^), and other Reactome pathways. Viral mRNAs in the cytoplasm can be translated by the host cell ribosomal translational apparatus, and indeed viruses have evolved strategies to recruit the host translation initiation factors necessary for the translation initiation by host cell mRNAs (*26*). NMD, a surveillance pathway that targets mRNAs with aberrant features for degradation, may interfere with the hijacking of the host translational machinery (*27*). In the brain, NMD, as a post-transcriptional mechanism, affects neural development, neural stem cell differentiation decisions, and synaptic plasticity, and thus defects in the pathway can cause aberrant neuronal activation and neurodevelopmental disorders (*18*). Detecting this coexpression network of ribosomal proteins therefore provides a sanity check to our approach, but the identified community structure and the presence of this in the multiplex may suggest a highly coordinated regulatory mechanism across the tissues.

Integration of the communities into TWAS of CRP in 361,194 subjects resulted in substantial performance gain in the discovery of trait-associated genes. Future methodological work that integrates community detection and additional omics data may further optimise the performance gain. A variational autoencoder implementation applied to the genes in the communities identified disease-relevant latent subspaces. Notably, this model learned a latent representation that significantly distinguishes metastastic from primary tumours and another related to stem cell-associated molecular features, across the 11,060 samples in TCGA data. Prediction of metastasis and the biology of the stemness phenotype can be further investigated through the genes and their communities identified here, but more definitive conclusions will require extensive biological and clinical validation studies. Nevertheless, methodologically, a deep learning framework that explicitly exploits the topological structures in co-expression networks holds promise for uncovering critical biological insights into disease mechanisms.

In summary, we performed network analysis on the most comprehensive human transcriptome dataset available to gain insights into how structures in co-expression networks may contribute to biological pathways and mediate disease processes. The rich resource we generated and the network approach we developed may prove useful to other omics datasets, facilitating studies of inter-tissue and intra-tissue regulatory mechanisms, with important implications for our mechanistic understanding of human disease.

## Materials and Methods

### GTEx Dataset

The GTEx V8 dataset (*7, 11*) is a genomic resource consisting of 948 donors and 17,382 RNA-Seq samples from 52 tissues and two cell lines. The resource provides a catalog of genetic effects on the transcriptome and a broad survey of individual- and tissue-specific gene expression. Of the 54 tissues and cell lines, 49 include samples with at least 70 subjects, forming the basis of the analysis of genetic regulatory effects (*7*). In this study, we leveraged the 49 tissues because of their sample size (see Table S1) and our interest in shared transcriptional regulatory programs for co-expressed genes.

### Data Preprocessing

We restricted our analyses to protein-encoding genes based on the GENCODE annotation. Although the GTEx dataset had annotated genes with ENSEMBLE IDs, we removed duplicates (using GENE IDs) and unmapped genes from downstream analyses. After this preprocessing step, the resulting dataset is characterised by the following count statistics:

- Unique genes across all tissues: 18,364
- Genes present in only 1 tissue: 412
- Genes present in all 49 tissues: 12,557

### Accounting for Unmodelled Factors

In order to correct for batch effects and other unwanted variation in the gene expression data, we used the *sva* R package, which is specifically targeted for identifying surrogate variables in high-dimensional data sets (*28*). For each tissue gene expression matrix, the number of components (latent factors) was estimated using a permutation procedure, as described in (*29*). Subsequently, using the function *sva network*, residuals were generated after regressing out the surrogate variables. The residual values, rather than the original gene expression values, were used in the downstream analyses. For convenience, we refer to the residual values as the ‘gene expression data’, since they represent the expression levels that have been corrected for (unwanted) confounders.

### Tissue-dependent Correlation and Adjacency Matrices

For each tissue, a correlation matrix *C* = [*z*_*ij*_] was created by calculating the Pearson correlation coefficient *r*_*ij*_ for every pair (*i, j*) of genes. Fisher *z*-transformation was then applied:

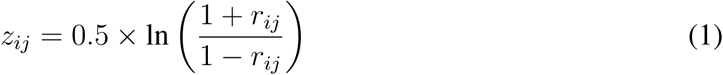

where *ln* is the natural logarithm function.

For each correlation matrix, we retained only the strongest correlations (i.e., transformed *z*_*ij*_ less than −0.8 and greater than 0.8) to generate a co-expression network. An adjacency matrix *A* = [*A*_*ij*_] was defined, for each tissue, such that *A*_*ij*_ is equal to *z*_*ij*_ if gene *i* and gene *j* are co-expressed (retained), and zero otherwise. We assumed undirected networks without self-loops, which implies *A*_*ij*_ = *A*_*ji*_ and *A*_*ii*_ = 0.

### Community Detection

We sought to detect groups of genes in each tissue, with the aim of finding communities whose internal connections are denser than the connections with the rest of the co-expression network. We applied the Louvain community detection method (*30*) in each tissue to generate a comprehensive atlas of communities. An asymmetric treatment for the negative correlations was used, thus inducing negatively correlated genes to belong to different communities (*31*). The algorithm identifies communities by maximising the modularity index (*32*), *Q*\*, as the algorithm progresses:

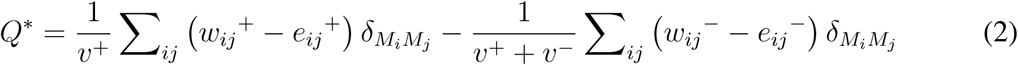

Here a positive connection between nodes *i* and *j* is denoted as *w*_*ij*_^+^ and has a value between 0 and 1; likewise, a negative connection is represented *w*_*ij*_^−^ and can also have a value between 0 and 1. *e*_*ij*_^±^ is the chance-expected within-module connection weight and calculated, for each positive/negative correspondent, as 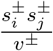, where 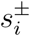 is the sum of positive or negative connection weights of node *i. v*^±^ is the sum of all positive or negative edges, and 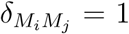 are in the same module or zero otherwise. In particular, the Louvain method iteratively evaluates the gain in modularity if one node is moved from one formed community to another of its neighborhood. We leveraged the Brain Connectivity Toolbox Python package (available on github: aestrivex/bctpy). The resolution parameter *γ* was set to its default value, 1.

### UMAP Embeddings of Community-defined Gene Expression

To produce a lower dimensional representation of the original dataset, we applied Uniform Manifold Approximation and Projection (UMAP) (*8*), a manifold learning technique. Our goal was to generate a map that reveals embedded structures and test whether biologically relevant clusters can be recovered from the gene expression data. Towards this end, we analysed both the full master matrix **M** of scaled gene expression (in the range [0, 1]), consisting of all genes (i.e., 18, 364), and a submatrix consisting of only those genes that belong to a community in at least one tissue (i.e., 3, 259). (Similarly to all of the results in the rest of the paper, we considered only Louvain communities with at least 4 genes.)

We chose UMAP because of the substantial improvement in running time on our data (compared to t-SNE, with its known computational and memory complexity that is quadratic in the sample size (*33*)), and UMAP’s theoretical grounding in manifold theory (*8*). UMAP can also capture non-linear effects in gene expression, and this was another reason why we chose it, over more traditional dimensionality reduction techniques such as Principal Component Analysis. Additional implementation details can be found in Figure S1.

### Persistence of the UMAP Global Structure

We quantified the conservation of, and variability in, the UMAP structure, including the relation among biologically-meaningful clusters, e.g., tissues. We characterised such a structure using the matrix [d(*i, j*)] of pairwise distances for clusters *i* and *j* in {1, 2, …, *L*}. For the actual (original) gene expression data, we define **V**_(**0**)_ = [*v*_*ij*,0_] as the resulting matrix of pairwise distances. Note **V**_(**0**)_ is a symmetric matrix with zeros along the diagonal. We sought to:

- estimate the sampling distribution of d(*i, j*), and calculate its standard error and a confidence interval
- correlate the matrix **V**_(**0**)_ and the resulting matrix *W* from a perturbation of the original structure

We approached the quantification problem through a (non-parametric) bootstrapping procedure. From the master matrix **M** of gene expression, we generated a total of *B* bootstrapped manifolds, each of equal size (here, each such sample was randomly drawn from 80% of the data points, i.e., rows, in **M**). For the *k*-th sample, we constructed the matrix 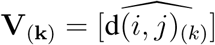 of pairwise distances derived from the UMAP embeddings for tissues *i* and *j*. Here we used the “induced metric” d : ℝ^*m*^ × ℝ^*m*^ → ℝ from the embedding *ϕ* : *M*_*g*_ → ℝ^*m*^ of the Riemannian manifold *M*_*g*_ into Euclidean space, but our treatment here generalises to the intrinsic metric g : *M*_*g*_ × *M*_*g*_ → ℝ, with g(*ϕ*^−1^(*i*), *ϕ*^−1^(*j*)), of the original manifold. The set 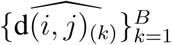 allows us to calculate the mean and variance of the UMAP-derived estimator for d(*i, j*):

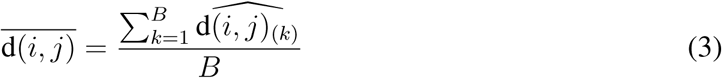

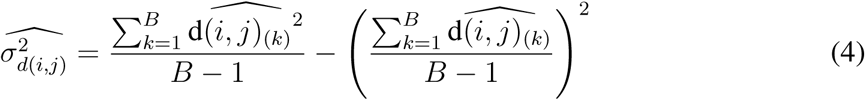

This approach provides a maximum likelihood estimate, i.e., 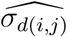, of the standard error. We used a heatmap to visualise 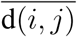 for each tissue pair (*i, j*). An alternative could have been to use a normalised “metric” (which is more robust to the scale from the embedding *ϕ*):

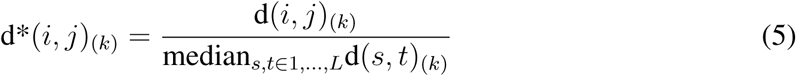

but we found this normalisation to be unnecessary in the GTEx data.

For two tissues *i*_0_ and *i*_1_, we define a “clustering conservation coefficient” to quantify the preservation of the clustering of tissues *i*_0_ and *i*_1_ relative to all tissues {*j*}:

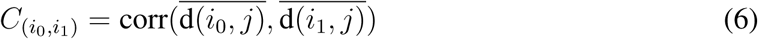

where *corr* is the correlation operator. In particular, this statistic allows us to formally test the null hypothesis of no conservation of global structure for a given pair of tissues; under the null hypothesis,

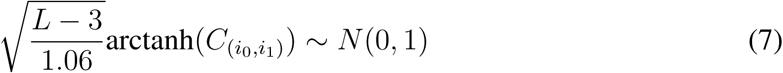

This coefficient can be extended to a larger set of tissues, *i*_0_, …, *i*_*l*_ (e.g., the 13 brain regions), using the first order statistic:

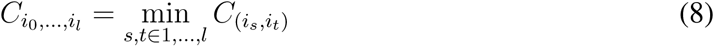

Furthermore, we calculated the relationship between the original **V**_(**0**)_ and “perturbed” **V**_(**k**)_ for each sample *k*:

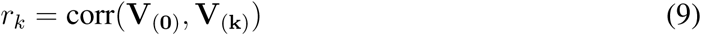

and the resulting empirical distribution of the correlation values *r*_*k*_. We note that UMAP has a stochastic element since it utilises stochastic approximate nearest neighbor search and stochastic gradient descent for optimisation; however, the *r*_*k*_ derived from a different run **V**_(**k**)_ (rather than from bootstrapping) quantifies the stability of the global structure in the presence of stochasticity. Collectively, our approach provides a way to perform statistical inference on the UMAP embedded structures.

### Prediction Power of Communities for Tissues

We investigated the extent to which each community’s gene expression profile was predictive of each of the tissues. The master matrix **M**, representing the entire dataset under analysis, has 15, 201 rows representing each RNA-Seq sample from each tissue collected from all subjects, and 18, 364 columns representing the total number of genes available. If a value was nonexistent (which may be due to the gene’s expression being tissue-specific), we assumed a zero value, conveying no expression in that tissue.

For each community, the expression values of the member genes were selected from **M**. With this sliced table, 49 binary classifications were performed using Support Vector Machine (SVM), wherein for each classification, we predicted each tissue. Essentially, the sliced table, which comprises the training data, for a *k*-member community can be viewed as a collection of vectors 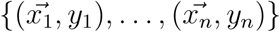, where 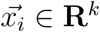 is the gene expression profile of the *k* genes for the *i*-th sample and *y*_*i*_ ∈ {1, 0} indicates membership in the tissue to be predicted. The goal of the classification is to separate the tissue to be predicted from the other tissues via the largest margin hyperplane, which can be generically written as 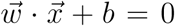, where 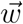 is normal to the hyperplane. SVM was used with a linear kernel and weights were adjusted to be inversely proportional to class frequencies in the input data (this corresponds to setting the *class weight* parameter in *scikit-learn* to “balanced”). To avoid overfitting, each classification was performed using a stratified 3-fold cross-validation procedure, in which the *F*_1_ score metric

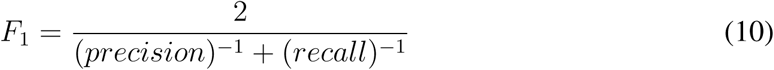

was used to report the prediction power across the three folds. We decided to use the *F*_1_ score instead of other metrics, given that each binary classification was highly unbalanced, i.e., a given tissue is the positive outcome and all the other 48 tissues are the negative outcome. (Louvain communities with less than 4 genes were filtered out from this analysis.)

### Enrichment Analysis

To evaluate the degree to which a community corresponds to well-known biological pathways, we performed enrichment analyses using the Reactome 2016 as reference. We used the *gseapy* python package to make calls on the *Enrichr* web API (*34*), and considered significant those pathways with a Benjamini-Hochberg-adjusted p-value below 0.05. (Louvain communities with less than 4 genes were considered “not enriched.”)

### Multilayer Analysis

In order to investigate the tissue-shared profiles of gene communities, as well as the relationships between gene expression traits across tissues, we proceeded to model our system as a multilayer network (*35*). Formally, a multilayer network is defined as a pair **Λ** = (**G**; **D**), where **G** := {*G*_1_, …, *G*_*L*_} is a set of graphs and **D** consists of a set of interlayer connections existing between the graphs and connecting the different layers. Each graph *l* ∈ **G** is a “network layer” with its own associated adjacency matrix *A*_*l*_. Thus, **G** can be specified by the vector of adjacency matrices of the *L* layers: **A** := (*A*_1_, …, *A*_*L*_). Multilayer networks allow us to represent complex relationships which would otherwise be impossible to describe using single-layer graphs separately considered.

A special case of multilayer networks is a multiplex network, which we used to model the GTEx transcriptome data. In this case, all layers are composed of the same set of nodes but may exhibit very different topologies. The degree of node *i* is the vector 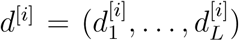, and 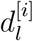 may vary across the layers. Interlayer connections are established between corresponding nodes across different layers. Layers represent different tissues, nodes represent genes, and edges between two nodes are weighted according to the correlation weights. In the GTEx data, the correlation matrices, previously described, define an adjacency matrix *A*_*l*_ for each layer *l* of the multiplex network. We applied community detection analysis to each layer separately to identify communities of co-expressed genes in each tissue, using the Louvain community detection method previously described.

Having identified communities of co-expressed genes for each tissue separately, we then computed the so-called *global multiplexity index* (*36*) to investigate the relationships of communities across different layers. This index quantifies how many times two nodes (genes) are clustered in the same communities across different layers. If, for example, gene *i* and gene *j* are clustered together in the layer of tissue *T*_1_ and of tissue *T*_2_, then the global multiplexity index is two. In the matrix [gmi(*i, j*)] of global multiplexity indices for a multiplex architecture, each element represents the number of times that two given genes, *i* and *j*, are clustered in the same community. More formally, if *L* is the number of layers, *g* is the generic layer, and *N* the number of nodes for each layer, then the global multiplexity index gmi(*i, j*) for gene *i* and gene *j*, with *i* and *j* ∈ {0, …, *N*} is defined as follows:

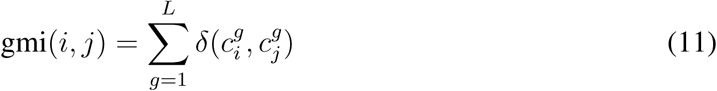

where 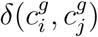 represents the Kroenecker delta function. The value of gmi(*i, j*) therefore increases by 1 if the two nodes are found to be part of the same community in a layer. If two genes share a high value of global multiplexity index, this may indicate a greater level of connectivity and suggest greater functional similarity, as they appear multiple times in the same community across different layers. We define the individual probabilities:

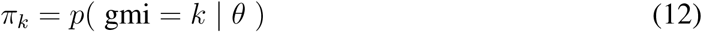

where *θ* represents all of the parameters of the model. *π*_*k*_ is the probability that two genes are clustered in the same communities across *k* layers, where *k* ≤ *L* and *L* is the number of layers in the network. We estimated the probability distribution of gmi(*i, j*) in the GTEx data.

We tested whether the UMAP embeddings of the transcriptome profiles of the communities in a multiplex architecture – a subset of all communities previously interrogated – could also recover biologically-meaningful clusters. This analysis allowed us to estimate the topology of the high-dimensional transcriptome data and test whether additional clusters could be uncovered at increasingly finer scales.

### Application to Transcriptome-Wide Association Studies

To evaluate the relevance of the communities for genomic studies of human disease, we performed TWAS / PrediXcan analysis of C-reactive protein (CRP) (*3*). We chose CRP for its clinical significance as a biomarker for a wide range of complex diseases, including cardiovascular disease, type 2 diabetes mellitus, Alzheimer’s disease, and age-related macular degeneration (*37*). Briefly, TWAS / PrediXcan estimates the “genetically-determined component of gene expression” in genome-wide association study (GWAS) subjects and infers the gene’s association with the phenotype. The inference can be done using GWAS summary statistics (*4, 14*). We hypothesised that integration of the communities into TWAS / PrediXcan would improve the signal-to-noise ratio for detecting gene-level associations. We applied whole blood models trained in GTEx v8 data (*38*) to a GWAS of CRP in 361,194 samples (of white-British ancestry) in the UK Biobank (*39*) (nealelab.is). We compared the associations derived from the set of genes that belong to a community and from the complement set of genes using a conditional Quantile-Quantile (Q-Q) plot (i.e., conditional on community membership status) of empirical quantiles of nominal negative log_10_(*p*) values. We estimated the true positive rate for each of the two non-overlapping sets of genes defined by community membership using the *π*_1_ statistic, as previously described (*14*).

### Variational Autoencoder Model of Communities

We implemented a variational autoencoder (VAE) (*23*), a deep learning methodology, to learn biologically-meaningful latent representations of the transcriptome (*5*) or a subset, e.g., the genes in the communities. VAE is a two-phase generative model, and we implemented it to capture major sources of variation with non-linear effects. In the encoding phase, dimensionality reduction is performed on the input; the decoding phase performs reconstruction of the original input from a latent and stochastic representation. The following equation serves as the basis for the VAE:

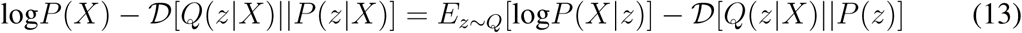

where *P* (*X*) is a probability density defined for a datapoint *X* in the input expression data (e.g., the set of genes that belong to the communities), *z* is a vector of latent variables with a prior probability density function *P* (*z*), and *D* is the Kullback-Leibler (KL) divergence between *P* (*z*|*X*) and some function *Q*(*z*). The first part of the right hand side of the equation, i.e., the expected negative log-likelihood, gives the reconstruction loss while the second is the KL divergence between the learned latent distribution and the prior distribution. *Q*(*z*|*X*) is the probabilistic encoder which compresses the data *X* into the latent variable *z* whereas the generative model *P* (*X*|*z*) is the probabilistic decoder which reconstructs the latent representation into the original data.

Using a VAE model of the communities, we tested the extent to which they could help to identify expression changes associated with disease and discover biologically-meaningful features (*5, 40*). We built on *Tybalt*, which uses a *Keras* implementation. We analysed TCGA Pan-Cancer data, consisting of batch-effects-normalised mRNA data (in units of log_2_(*norm value*+1)) in 11,060 samples across 33 cancer types (see “Data and materials availability”). Expression values were mapped to [0, 1] for each gene using the maximum and minimum values.

We evaluated the performance of the VAE model of the GTEx-derived communities in TCGA data in two ways. (1) The process of metastasis remains poorly understood. Classification of the tumours into primary or metastatic origin enabled us to test whether the VAE model could successfully discriminate the sample type. (2) The acquisition of a stem cell-like tumour trait, i.e., stemness, in cancer suggests gene expression programs that may contribute to progression and treatment resistance. To determine whether the VAE model successfully learned stemness (*24*), we used a DNA methylation-based “Stemness Score” from the Pan-Cancer Stemness working group, specifically the “DNAss” signature, which combines (a) an epigenetically-regulated DNA methylation-based signature, (b) a differentially-methylated probes-based signature, and (c) an enhancer elements / DNA methylation-based signature.

## Acknowledgements

E.R.G. is grateful to Clare Hall, University of Cambridge for the Fellowship support. We thank The Genotype-Tissue Expression (GTEx) project for the use of v8 data and The Cancer Genome Atlas (TCGA) project for the use of gene expression data from the cancer samples. This research has been conducted using the UK Biobank Resource.

## Funding

This research is supported by the National Institutes of Health Genomic Innovator Award (NHGRI R35HG010718). T.A. is funded by the W. D. Armstrong Trust Fund, University of Cambridge, UK.

## Author contributions

T.A. and G.M.D. conducted the analysis. E.R.G, T.A., G.M.D. wrote the manuscript. E.R.G. and P.L. designed and supervised the study. All authors contributed to the editing of the manuscript.

## Competing interests

E.R.G. receives an honorarium from the journal Circulation Research of the American Heart Association, as a member of the Editorial Board. He performed consulting on pharmacogenetic analysis with the City of Hope / Beckman Research Institute. The other authors declare no competing interests.

## Data and materials availability

The protected data for the GTEx project (for example, genotype and RNA-sequence data) are available via access request to dbGaP accession number phs000424.v8.p2. Processed GTEx data (for example, gene expression and eQTLs) are available on the GTEx portal: https://gtexportal.org. TCGA gene expression data and DNA methylation based stemness scores can be downloaded from the University of California, Santa Cruz (UCSC) TCGA Pan-Cancer Atlas hub on Xena: https://pancanatlas.xenahubs.net. The UK Biobank C-Reactive Protein GWAS summary results on which TWAS / PrediXcan was applied are available for download: https://www.dropbox.com/s/4o9qs1s6jkrzrq7/30710_irnt.gwas.imputed_v3.both_sexes.tsv.bgz?dl=0-O30710_irnt.gwas.imputed_v3.both_sexes.tsv.bgz. Results and code (for reproducibility) are available on github: https://github.com/tjiagoM/gtex-transcriptome-modelling.

## Notes

### Summary of Updates

paragraph reordering. sentences about confounds correction on previous experiments

