## Supplementary Material for "Multilayer modelling and analysis of the human transcriptome"

#### Methods

**Prediction of Tissues Given Reactome Pathways.** For comparison with the communities, we investigated the extent to which biologically meaningful sets of genes encoding current biological knowledge are predictive of tissues. For each Reactome (*1*) pathway, we selected the expression of member genes from the master matrix M. If a gene from a Reactome pathway was not present, that gene was ignored. Using the same stratified 3-fold cross validation procedure described in Methods of the main text, we performed 49 binary classifications.

**Embedding new transcriptome data into UMAP learned space.** To evaluate the relevance of the trained model generated from the GTEx communities, we passed previously unseen data $D_{test}$ to the model for embedding into the learned latent map (from the UMAP embedding of GTEx training data, $\phi : D_{train} \subset M_g \hookrightarrow \mathbb{R}^m$). We used The Cancer Genome Atlas (TCGA) gene expression data in acute myeloid leukemia (*2*), breast cancer (*3*), and lung adenocarcinoma (*4*) as test sets.

### Results

**Relationship between Reactome Pathways and Tissues.** Consistent with our observations for the communities, most Reactome pathways are not sufficient to predict any tissue (available on github: output *output_06_02*), while many are tissue-specific (i.e., can predict only one tissue). However, we identified Reactome pathways that can predict more than half of the tissues: *GPCR LIGAND BINDING*, *GPCR DOWNSTREAM SIGNALING*, and *SIGNALING BY GPCR* predict 34, 33, and 32 tissues, respectively. This observation is perhaps expected: G-protein-coupled receptors (GPCRs) comprise a large family of cell surface receptors that form the essential sites of communication between the internal and external environments of cells, with a central and widespread role in human physiology (*5*). Their gene expression profile in each of the predicted tissues differs from the remaining tissues, potentially reflecting their broad but tissue-specific function.

Prediction of tissues by Reactome pathways varies substantially. The brain tissues “Brain Caudate” (basal ganglia), “Brain Frontal Cortex”, “Brain Hippocampus”, and “Brain Nucleus” accumbens (basal ganglia) are not predicted by any Reactome pathway, likely reflecting the fact that our current understandings (as encoded in these pathways) have been hampered by the relative inaccessibility of these tissues. In contrast, the tissues cells cultured fibroblasts and whole blood are the tissues most highly predicted (269 and 330 Reactomes, respectively). Some tissues are predicted by less than 5 Reactome pathways, including the tissues “Brain Amygdala” (two), “Brain Anterior Cingulate” cortex (BA24) (one), “Brain Cortex” (two), and “Brain Hypothalamus” (two). More information on the relationship between communities and enriched pathways is available on github: notebook *10_reactomes_per_tissue*.

**Embedding new transcriptome data into UMAP learned space.** Interestingly, embedding of each of the TCGA datasets into the learned latent space from the communities showed clustering with the testis tissue. This result recapitulates two known results: (a) the GTEx finding that the testis is an outlier relative to the other GTEx tissues in transcriptome profile (*6*) and (b) the role of the so-called cancer-testis (CT) genes (*7*), which function as driver genes in cancer (*8, 9*), evoke immune responses in cancer patients as immunogenic antigens across a range of cancers (*10*), and contribute to various neoplastic phenotypes. The implementation is available on github: notebook *12_tcga*.

### Supplementary Figures

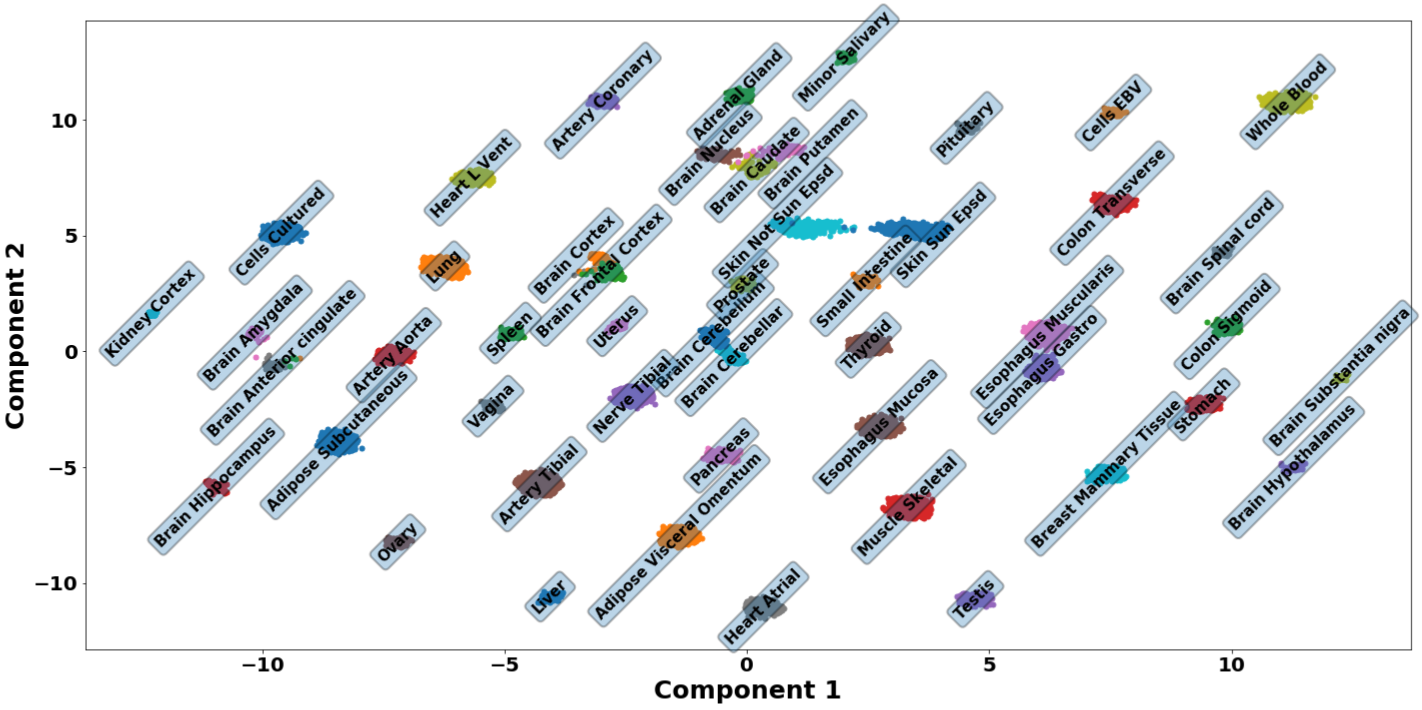


Figure S1: **Lower-dimensional representation of the transcriptome data.** The plot shows the UMAP embedded components from the full master matrix consisting of all genes. The clusters formed in the embedding space are well-defined. The UMAP parameters used in this study to create the reduced embeddings assumed the default values except for the the size of local neighbourhood (i.e., $5$), the effective minimum distance between embedded points (i.e., $0.25$), the learning rate (i.e., $0.5$), and the spread (i.e., $1$).

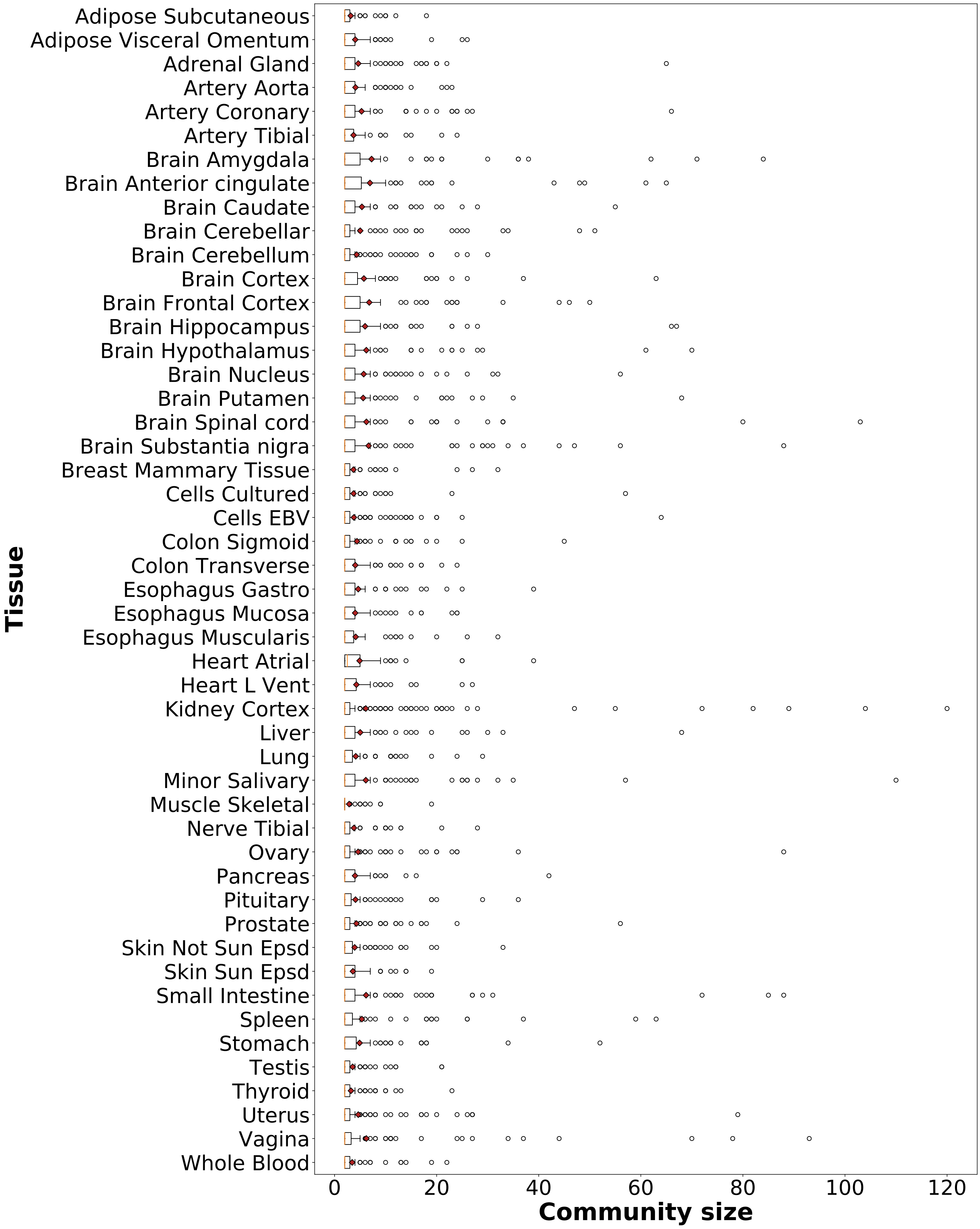


Figure S2: **Distribution of community size for each tissue.** The boxplots show the distribution of community size for each of the 49 tissues. Community size varies within each tissue and across tissues.

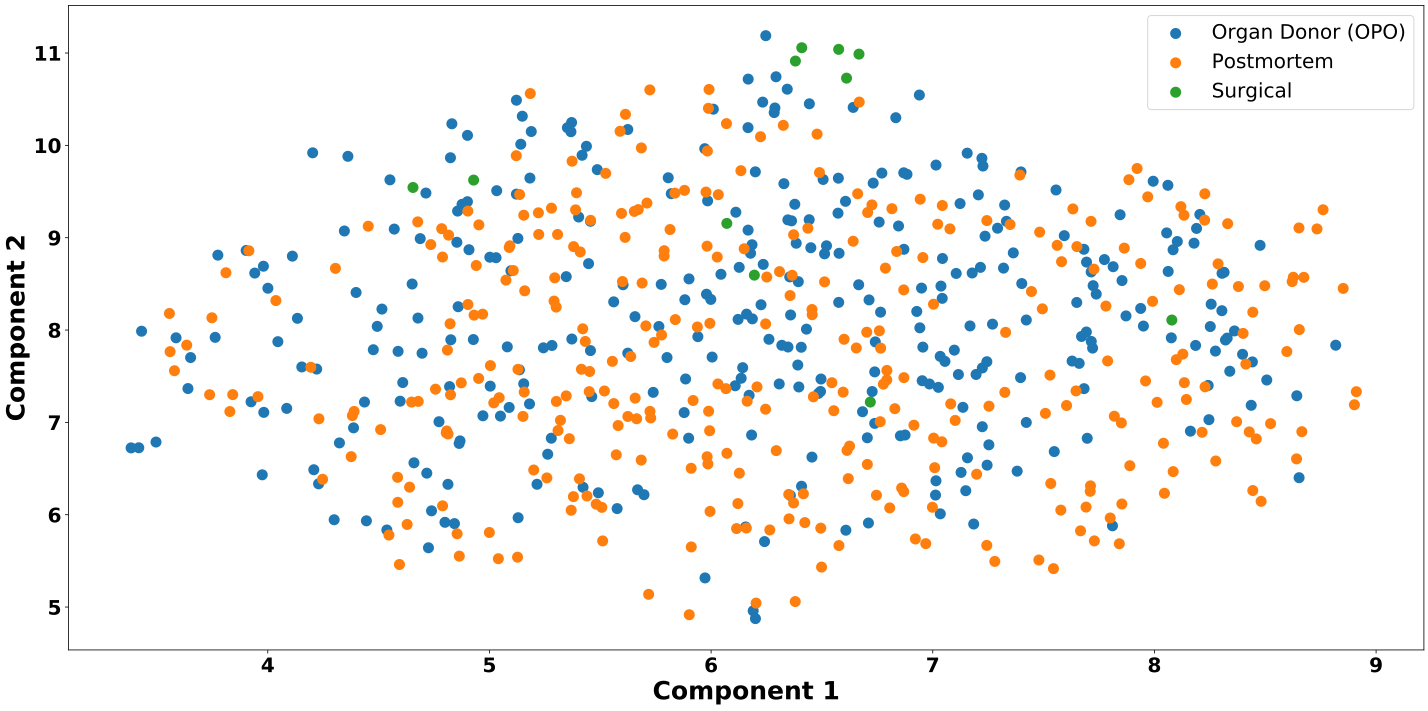


Figure S3: **Lower-dimensional representation of the whole blood transcriptome, overlayed with COHORT values.** The UMAP embedding shows no clustering in the whole blood structure with regards to COHORT, indicating that this confound was successfully corrected with *sva*.

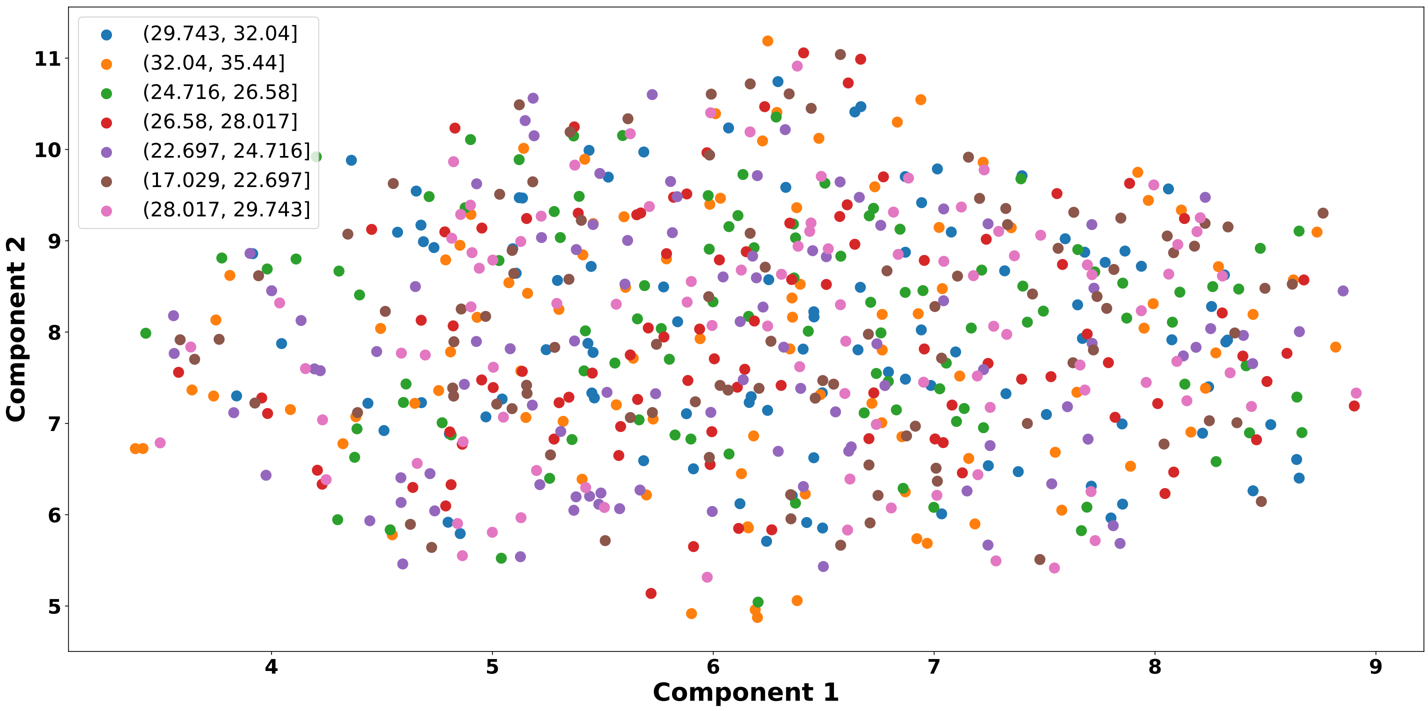


Figure S4: **Lower-dimensional representation of the whole blood transcriptome, overlayed with BMI values.** The UMAP embedding shows no clustering in the whole blood structure with regards to BMI, indicating that this confound was successfully corrected with *sva*.

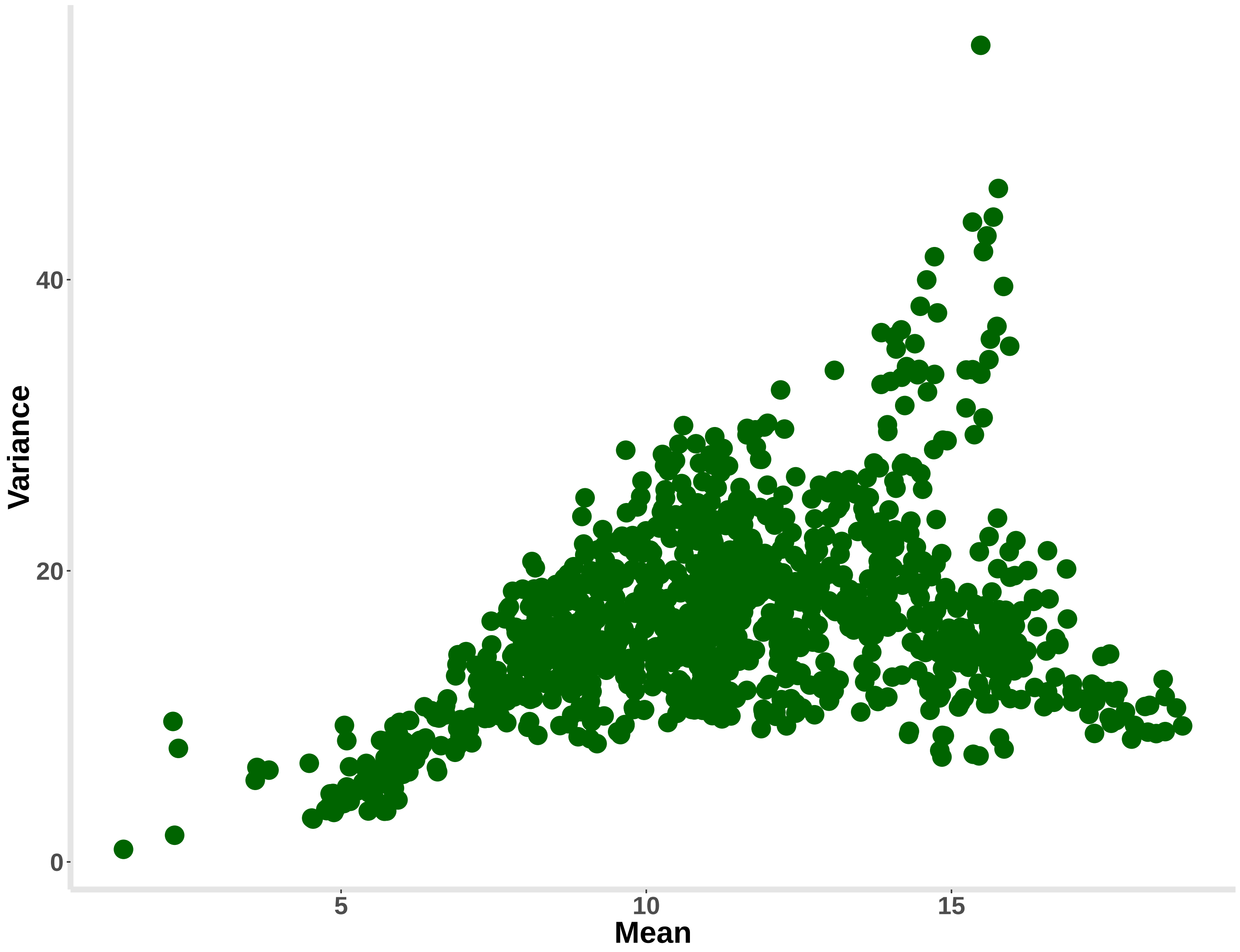


Figure S5: **Mean and variance of distance between clusters across bootstrapped manifolds.** There is a significant correlation between mean and variance (Spearman $\rho \approx 0.38$, $p < 2.2 \times 10^{-16}$) across bootstraps. For a pair of tissue clusters, greater average distance implies greater variability in the distance.

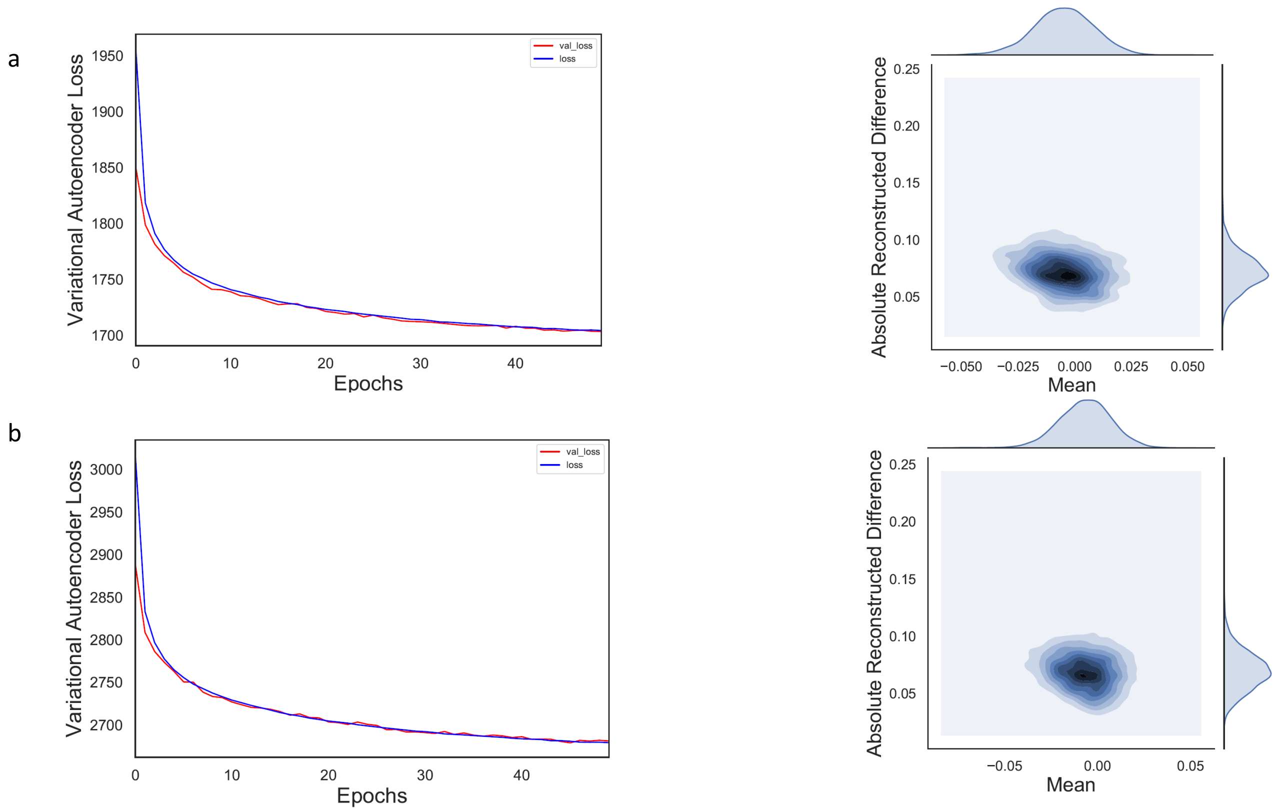


Figure S6: **Diagnostics.** **a.** The training performance (VAE loss as a function of epoch number) and reconstruction accuracy (mean vs absolute difference) of the VAE applied to the genes in the communities are shown (top panel). **b.** The corresponding plots for the full set of genes regardless of community membership are shown (bottom panel). *Tybalt* code was modified to generate these diagnostic plots.

### Supplementary Tables

Table S1: **Sample size of each tissue in GTEx V8.**

| Tissue | Sample Size |
|---|---|
| Adipose Subcutaneous | 581 |
| Adipose Visceral Omentum | 469 |
| Adrenal Gland | 233 |
| Artery Aorta | 387 |
| Artery Coronary | 213 |
| Artery Tibial | 584 |
| Brain Amygdala | 129 |
| Brain Anterior cingulate | 147 |
| Brain Caudate | 194 |
| Brain Cerebellar | 175 |
| Brain Cerebellum | 209 |
| Brain Cortex | 205 |
| Brain Frontal Cortex | 175 |
| Brain Hippocampus | 165 |
| Brain Hypothalamus | 170 |
| Brain Nucleus | 202 |
| Brain Putamen | 170 |
| Brain Spinal cord | 126 |
| Brain Substantia nigra | 114 |
| Breast Mammary Tissue | 396 |
| Cells Cultured | 483 |
| Cells EBV | 147 |
| Colon Sigmoid | 318 |
| Colon Transverse | 368 |
| Esophagus Gastro | 330 |
| Esophagus Mucosa | 497 |
| Esophagus Muscularis | 465 |
| Heart Atrial | 372 |
| Heart L Vent | 386 |
| Kidney Cortex | 73 |
| Liver | 208 |
| Lung | 515 |
| Minor Salivary | 144 |
| Muscle Skeletal | 706 |
| Nerve Tibial | 532 |

Continued on next page

Table S1: **Sample size of each tissue in GTEx V8.**

| Tissue | Sample Size |
|---|---|
| Ovary | 167 |
| Pancreas | 305 |
| Pituitary | 237 |
| Prostate | 221 |
| Skin Not Sun Epsd | 517 |
| Skin Sun Epsd | 605 |
| Small Intestine | 174 |
| Spleen | 227 |
| Stomach | 324 |
| Testis | 322 |
| Thyroid | 574 |
| Uterus | 129 |
| Vagina | 141 |
| Whole Blood | 670 |

Table S2: **Information about communities in each tissue.** Statistics (i.e., mean, std, median, and MAD) refer to the distribution of community size for each tissue

| Tissue | Number of genes without a community | Number of communities with size below 4 | Total number of communities | Mean | Std | Median | MAD |
|---|---|---|---|---|---|---|---|
| Adipose Subcutaneous | 15770 | 69 | 84 | 3.167 | 2.6763 | 2.0 | 0.0 |
| Adipose Visceral Omentum | 15730 | 62 | 91 | 4.055 | 4.2462 | 2.0 | 0.0 |
| Adrenal Gland | 15260 | 93 | 125 | 4.648 | 7.0373 | 2.0 | 0.0 |
| Artery Aorta | 15463 | 73 | 100 | 4.1 | 4.2837 | 2.0 | 0.0 |
| Artery Coronary | 15490 | 80 | 112 | 5.286 | 8.2163 | 2.0 | 0.0 |
| Artery Tibial | 15402 | 58 | 78 | 3.718 | 3.9545 | 2.0 | 0.0 |
| Brain Amygdala | 15430 | 71 | 102 | 7.255 | 13.5536 | 2.0 | 0.0 |
| Brain Anterior cingulate | 15499 | 67 | 96 | 6.938 | 11.868 | 2.0 | 0.0 |
| Brain Caudate | 15748 | 57 | 89 | 5.36 | 7.5032 | 2.0 | 0.0 |
| Brain Cerebellar | 15402 | 106 | 136 | 5.029 | 8.0229 | 2.0 | 0.0 |
| Brain Cerebellum | 15713 | 88 | 113 | 4.327 | 5.2131 | 2.0 | 0.0 |
| Brain Cortex | 15765 | 60 | 91 | 5.747 | 8.6642 | 2.0 | 0.0 |
| Brain Frontal Cortex | 15638 | 58 | 89 | 6.798 | 9.861 | 2.0 | 0.0 |
| Brain Hippocampus | 15642 | 67 | 103 | 5.981 | 10.0242 | 2.0 | 0.0 |
| Brain Hypothalamus | 15885 | 66 | 94 | 6.223 | 10.6553 | 2.0 | 0.0 |
| Brain Nucleus | 15725 | 63 | 86 | 5.709 | 8.3525 | 2.0 | 0.0 |
| Brain Putamen | 15470 | 70 | 96 | 5.604 | 9.0559 | 2.0 | 0.0 |
| Brain Spinal cord | 15550 | 93 | 129 | 6.24 | 12.7788 | 2.0 | 0.0 |

Continued on next page

Table S2: **Information about communities in each tissue.** Statistics (i.e., mean, std, median, and MAD) refer to the distribution of community size for each tissue

| Tissue | Number of genes without a community | Number of communities with size below 4 | Total number of communities | Mean | Std | Median | MAD |
|---|---|---|---|---|---|---|---|
| Brain Substantia nigra | 15241 | 99 | 139 | 6.669 | 11.8099 | 2.0 | 0.0 |
| Breast Mammary Tissue | 16050 | 77 | 98 | 3.704 | 4.7385 | 2.0 | 0.0 |
| Cells Cultured | 14525 | 75 | 94 | 3.723 | 6.1995 | 2.0 | 0.0 |
| Cells EBV | 14052 | 151 | 185 | 3.784 | 5.7673 | 2.0 | 0.0 |
| Colon Sigmoid | 15811 | 70 | 92 | 4.391 | 6.0666 | 2.0 | 0.0 |
| Colon Transverse | 15946 | 83 | 116 | 4.06 | 4.1028 | 2.0 | 0.0 |
| Esophagus Gastro | 15609 | 60 | 86 | 4.64 | 5.8387 | 2.0 | 0.0 |
| Esophagus Mucosa | 15556 | 77 | 105 | 4.038 | 4.4953 | 2.0 | 0.0 |
| Esophagus Muscularis | 15573 | 64 | 86 | 4.14 | 5.0513 | 2.0 | 0.0 |
| Heart Atrial | 15265 | 51 | 76 | 4.908 | 5.9564 | 2.5 | 0.5 |
| Heart L Vent | 14654 | 51 | 76 | 4.276 | 4.6043 | 2.0 | 0.0 |
| Kidney Cortex | 14667 | 192 | 251 | 6.116 | 14.1071 | 2.0 | 0.0 |
| Liver | 14769 | 80 | 116 | 5.026 | 8.0456 | 2.0 | 0.0 |
| Lung | 16090 | 74 | 99 | 4.152 | 4.5911 | 2.0 | 0.0 |
| Minor Salivary | 15545 | 97 | 137 | 6.124 | 11.8639 | 2.0 | 0.0 |
| Muscle Skeletal | 14746 | 63 | 73 | 2.863 | 2.4231 | 2.0 | 0.0 |
| Nerve Tibial | 15975 | 66 | 84 | 3.798 | 4.228 | 2.0 | 0.0 |
| Ovary | 15478 | 109 | 142 | 4.62 | 8.6917 | 2.0 | 0.0 |
| Pancreas | 15297 | 59 | 80 | 4.012 | 5.0855 | 2.0 | 0.0 |
| Pituitary | 16318 | 81 | 108 | 4.102 | 5.2651 | 2.0 | 0.0 |
| Prostate | 16149 | 87 | 115 | 4.287 | 6.2147 | 2.0 | 0.0 |
| Skin Not Sun Epsd | 16023 | 74 | 99 | 3.939 | 4.559 | 2.0 | 0.0 |
| Skin Sun Epsd | 16082 | 64 | 87 | 3.598 | 3.12 | 2.0 | 0.0 |
| Small Intestine | 15680 | 101 | 145 | 6.172 | 12.3027 | 2.0 | 0.0 |
| Spleen | 15455 | 83 | 111 | 5.342 | 9.4654 | 2.0 | 0.0 |
| Stomach | 15767 | 67 | 96 | 4.906 | 6.9463 | 2.0 | 0.0 |
| Testis | 17679 | 70 | 91 | 3.538 | 3.3946 | 2.0 | 0.0 |
| Thyroid | 15994 | 93 | 111 | 3.216 | 2.8582 | 2.0 | 0.0 |
| Uterus | 15473 | 113 | 148 | 4.649 | 8.0395 | 2.0 | 0.0 |
| Vagina | 15596 | 99 | 132 | 6.227 | 13.1912 | 2.0 | 0.0 |
| Whole Blood | 14061 | 69 | 85 | 3.447 | 3.5695 | 2.0 | 0.0 |
